# Anxiety state-related task disengagement varies with trait anxiety

**DOI:** 10.1101/2025.05.04.651621

**Authors:** Ceyda Sayalı, Emma Heling, Roshan Cools

**Author notes:** Co-first authorship (Ceyda Sayalı, Emma Heling). <u>Corresponding Author:</u> Ceyda Sayalı, 5510 Nathan Shock Dr, Baltimore, MD 21224. **Author contributions:** CS: conceptualization, methodology, software, project administration, supervision, writing-original draft; EH: formal analysis, investigation, visualization, writing-original draft; RC: conceptualization, funding acquisition, supervision, writing-review and editing.

## Abstract

Cognitively demanding tasks are often perceived as costly due to the cognitive control resources they require, leading to effort avoidance, particularly in psychiatric populations with motivational impairments. Research on anxiety and cognitive effort are mixed: some studies suggest anxiety increases the perceived effort cost and avoidance, while others indicate that cognitive effort engagement can serve as an adaptive coping strategy. To reconcile these perspectives, we examined the interaction between state and trait anxiety on cognitive effort evaluation and engagement in two experiments. We hypothesized that state anxiety enhances task engagement as difficulty increases, and that this effect is diminished in individuals with high trait anxiety. Experiment 1 assessed self-reported anxiety in an online sample, while Experiment 2 manipulated state anxiety through autobiographical recall. Both experiments employed flow induction and effort discounting paradigms. Across both studies, the effect of state anxiety on task engagement depended on trait anxiety, but the direction of the state anxiety effect was opposite to the effect we predicted. In Experiment 1, participants with low trait anxiety reported reduced task engagement, as indexed by lower flow scores, when state anxiety was higher, but only in easy tasks. This effect was attenuated in participants with higher trait anxiety. The same pattern was observed in Experiment 2, but this time the interaction between trait and state anxiety was present regardless of task difficulty. These findings suggest that trait anxiety may reflect reduced impact of state anxiety on task disengagement.

**Public significance statement:** This study demonstrated that effects of state anxiety on task disengagement depend on individual differences in trait anxiety. People with higher trait anxiety reported reduced effects of state anxiety on task disengagement.

## Introduction

A growing body of literature suggests that people avoid cognitive effort when given the option (Kool et al., 2010; Sayali & Badre, 2019). This tendency to avoid mentally effortful activities rises to pathological levels in psychiatric populations marked with motivational problems (Le Heron, Apps, Husain, 2017). One prominent account of cognitive effort avoidance is the Expected Value of Control theory (Shenhav et al., 2013). This theory posits that cognitively effortful tasks are costly to the extent they recruit cognitive control resources. For example, it was shown that people who recruited greater behavioral and neural proxies of cognitive control for otherwise comparable task performance showed greater rates of effort avoidance (McGuire et al., 2010; Sayali & Badre, 2019), indicating that information processing capacity might mediate effort evaluation (Shenhav et al., 2017). As such, information processing capacity limitations have been theorized to underlie the transdiagnostic structure of mental effort avoidance (Gillan et al., 2016). Given that anxiety disrupts processing efficiency and increases the need for cognitive control resources to maintain task performance (Eysenck & Calvo, 1992; Shenhav et al., 2013), it follows that effortful tasks should be perceived as more costly under anxious states. Likewise, anxious-depression has been shown to correlate positively with demand avoidance in a demand selection task (Patzelt et al., 2019).

On the other hand, it has been suggested that cognitively effortful tasks might operate as an adaptive cognitive coping strategy by means of self-distraction from the source of anxiety (Kalisch et al., 2006). Accordingly, anxiety-related mental contents can be replaced by anxiolytic task-related thoughts that mentally challenging tasks offer. Therefore, one way to combat negative internal processes such as negative thoughts or ruminations, might be via attending to impersonal or task-related thoughts (Balderston et al., 2016). Indeed, Vytal and colleagues (2012) demonstrate that participants felt less electric-shock related anxiety (in terms of the startle reflex) when they were performing a 3-back working memory task compared with an easier 1-back working memory task, suggesting that the cognitive effort load might compete with mental resources necessary to entertain worrisome thoughts. Consistent with this interpretation, the same study showed that state anxiety impaired working memory performance only when load was low and not when it was high, suggesting that performing a challenging task had a protective effect on executive functions during a state of anxiety.

These two separate lines of research raise two conflicting hypotheses regarding the effects of anxiety on effort evaluation. These seemingly contradictory results can be reconciled in one unified account by taking into consideration the differential roles of state and trait anxiety. *State anxiety* is a transient state of arousal and is associated with a temporary increased sympathetic nervous system activity. Contrarily, *trait anxiety* is a personality trait that is associated with a psychopathological condition of constant high arousal. State anxiety can be induced by experimental conditions in the lab, whereas trait anxiety is a more stable characteristic of an individual. Although it is currently debated whether these two measures are different sides of the same coin or separate multidimensional constructs, recent research is beginning to suggest that state and trait anxiety might implicate distinct neuroanatomical and functional mechanisms (Saviola et al., 2020). While state anxiety has been shown to promote adaptive coping strategies in otherwise healthy individuals (Grillon, 2008), trait anxiety has been associated with maladaptive coping with environmental challenges and hyper-responsivity to stress (Weger & Sandi, 2018). Therefore, the effects of state anxiety might interact with trait anxiety so that trait anxiety might undermine the capacity of adopting adaptive coping strategies in the face of state anxiety.

Allostatic accounts of stress coping (e.g., McEwen & Wingfield, 2003) suggest that the cumulative burden of stress (allostatic load) may shape how individuals regulate physiological and psychological responses to new stressors. Trait anxiety could represent a form of chronic allostatic load, impairing the ability to engage in adaptive short-term (state-dependent) coping mechanisms. This aligns with the idea that while state anxiety can mobilize adaptive responses in healthy individuals, those with high trait anxiety may already be operating under a chronically dysregulated stress system, limiting their capacity to effectively deploy such adaptive strategies.

We hypothesized that people with greater levels of state anxiety experience increasingly effortful and challenging tasks as more engaging and enjoyable. However, this effect was expected to be diminished in participants with high trait anxiety. In these individuals, adaptive coping strategies might not be effectively recruited. This hypothesis predicts an interaction between state and trait anxiety on cognitive effort evaluation as a function of task difficulty. Specifically, we predicted that people with low trait anxiety exhibit a positive association between state anxiety and effort value as a function of increasing task difficulty. Conversely, this positive association between state anxiety and difficulty-dependent effort value was anticipated to be diminished in people with high trait anxiety.

The current study tests this hypothesis in an online sample using a flow induction paradigm that we previously developed (Sayali, Heling, Cools, 2022). In this paradigm, participants perform a task that consists of different levels of cognitive effort investment and are asked to evaluate their effort investments per level in terms of their experience of ‘flow’ (Csikszentmihalyi, 1990). Flow can be described as a state of mind in which a person experiences full engagement, involvement and enjoyment in an activity. This state occurs for activities with an intermediate challenge and skill level. Activities that are too easy have been established to elicit disengagement due to boredom, while activities that are too difficult result in disengagement due to anxiety and stress (Agrawal et al., 2022; Csikszentmihalyi, 1990). We also asked participants to evaluate cognitive effort in terms of the minimum monetary payoff they wanted to receive in return for exerting it (Shenhav, Botvenick, & Cohen, 2013; Westbrook, Kester, & Braver, 2013).

In the first experiment, we assess effects of self-reported state and trait anxiety on task engagement, as indexed by effortful task performance, and self-reported flow and effort value ratings. In the second experiment, we induce state anxiety by asking participants to write an essay about a personally significant, anxiety-producing event. This allowed us to assess the effects of induced state anxiety on task engagement as a function of self-reported trait and state anxiety as well as task difficulty.

We predicted a positive association between state anxiety and task engagement with increasing task difficulty in individuals with low trait anxiety, whereas this association would be attenuated in those with higher trait anxiety. The findings align with our hypothesis that the effects of state anxiety on task engagement are attenuated in people with higher trait anxiety. However, the direction of the effect of state anxiety in people with low trait anxiety was opposite to that predicted. While we had predicted a positive effect of state anxiety on task engagement as task difficulty increased, in fact, we observed a negative effect of state anxiety: Task engagement was reduced rather than enhanced with state anxiety. The finding in experiment 1 that this interaction between state and trait anxiety surfaced only when the task was overly easy raises the question whether trait anxiety attenuates the impact of anxiety state adaptation to low task difficulty, i.e. boredom-related disengagement (Agrawal et al., 2022).

## 2. Methods experiment 1

### 2.1 Participants

The initial sample included 126 English-speaking participants, recruited from the international online portal Prolific. Having participated in earlier versions of this experiment was considered an exclusion criterion. Prior to the experiment, all participants signed an online written informed consent to take part in the study, which was approved by the regional research ethics committee (Commissie Mensgebonden Onderzoek, Arnhem/Nijmegen, CMO2014/288). Each participant took part in an online session lasting approximately 80 minutes. All received a monetary reward (€8.66) for their participation with a bonus (€0.00-€1.00) depending on their choices. Participants were excluded from the analyses if they had not completed the study (N = 9) or if they scored 3 SD away on the average accuracy and RT in each of the four task levels (N = 10). The final sample consisted of 107 participants (aged 18-69; *M* = 29.79; *SD* = 12.11; 36 women).

### 2.2 Stimulus Presentation and Data Acquisition

The participants performed the experiment at home at their own laptop. Participants were directed from the recruitment portal to the experiment programmed, which was programmed using Gorilla Experimenter Builder. Stimuli consisted of arithmetic summations of diverse difficulty levels, manipulated by summation length. Experimental scripts used to create the task can be found on the author’s personal Github page (https://github.com/zceydas/OptEft).

### 2.3 Procedure

#### 2.3.1 Psychological inventory

Prior to the task, participants were asked to complete several inventories, among which the State-Trait Anxiety Inventory (STAI) which assesses self-reports of state and trait anxiety (Spielberger, 1983) (see supplementary section 8.1 and 8.2 for an overview of the inventories). Other inventories included were the Need for Cognition (NfC), Positive and Negative Affect Schedule (PANAS), and the Brief State Rumination Inventory (BSRI).

#### 2.3.2 The flow induction paradigm

Participants completed a computerized task consisting of two phases in which they solved arithmetic summations varying in difficulty. A detailed description of the task can be found in Sayali, Heling, Cools, 2023. In summary, participants had to solve summations by entering digits on their keyboard and were all provided with accurate feedback (*correct, incorrect*, or *too slow, presented for* 3 seconds) in both phases. The ‘capacity phase’ was implemented to determine four individually tailored difficulty levels. Starting at the easiest difficulty level, the difficulty level increased in a stepwise order of 5 summation questions until a score of 0% was reached (5 incorrect summations). Subsequently, four difficulty levels were determined by fitting a sigmoid function, scoring 100, 75, 50, and 25% correct (easy, intermediate1 intermediate2, difficult respectively). In the ‘performance phase’ participants performed each of these four difficulty levels once in a pseudo-randomized order (as opposed to two blocks of each difficulty level the original study). Each level consisted of 8 trials. Also, each trial started with an indicator of the upcoming difficulty level in the form of a color cue (*saddle brown, green, blue*, and *red*) that was counterbalanced across participants.

Subjective flow experiences were assessed by nine visual analogue items at the end of each level (Ulrich et al, 2014; Keller and Bless, 2008; Sayali, Heling, Cools, 2023). To do so, participants could indicate their response by moving the mouse on a horizontal line without any anchors, except for start (0) middle (50) and endpoint (100). The endpoints were labelled *agree* and *disagree*.

#### 2.3.3 Effort cost question

Participants completed four effort cost (EC) questions to assess their willingness to perform each difficulty level, at the end of the performance phase. This method is adopted from Becker, DeGroot, and Marschak (1964) and is used to provide an estimate of the subjective utility of money by measuring the cash-equivalent of a wager in a short sequential procedure. Such a procedure is ideal for an online sample compared with a longer effort cost procedure based on repeated decision making (Westbrook, Kester, and Braver, 2013).

Specifically, participants were asked to give a bid for each difficulty level on a scale ranging from 0.0 to 1.0 euro (steps of .05) to indicate the minimum amount of money that they would request in exchange for repeating a certain task difficulty level (later referred to as effort cost score). They were told that a random number would be drawn between 0.0 and 1.0 and that they had to repeat the difficulty level if that number was greater than their number for that level. If that number was equal or smaller, they would not repeat that difficulty level and earn nothing. Unknown to the participants, they had to repeat one of the difficulty levels regardless of their bids, which was chosen randomly. Their bid associated to that specific difficulty level was added to their payment as bonus. The full description of the instruction is provided in the supplementary method section 8.1.2.

### 2.4 Statistical analyses

All statistical analyses were conducted in Rstudio (Version 1.3.1093). Data were analyzed with analysis of variance (ANOVA) and linear mixed models (LMM). For the ANOVA, the R package *ez*. was used, and for the LMM models we used the lme4 package (Bates, Mächler, Bolker, & Walker, 2015).

Potential nuisance variables, including age, gender, years of education, and task order (order of difficulty condition presented in the task), were initially tested for their relationship with the dependent variables using Pearson correlation. Only variables with significant correlations (*p* > .05) were included as covariates in the models and are only reported when having a significant effect. All significant *(p* < .05) covariates are reported. Additionally, in all models, subject number was entered as random effect, and random slopes for difficulty level were included, providing the model did not exhibit singularity issues. Lastly, prior to each model fitting, predictor multicollinearity was tested by computing the variance inflation factor (VIF) index. This index measures the inflation of a regression coefficient due to collinearity between predictors and exceeding the threshold of 5 is considered problematic (Bruce & Bruce, 2017; James et al., 2014). Consequently, predictors that exceed this threshold are considered redundant. In this dataset, there was no multicollinearity between predictors (see supplementary method section 8.1, table 2). VIF scores and (relevant) model estimates of all four subjective flow subcomponents (involvement, liking, ability, time) can be found in supplementary section 8.1, table 4, 8.2, table 5-6).

#### 2.4.1 Correlation inventories

To investigate correlations between STAI-state and STAI-trait, Pearson’s tests were conducted to measure the linear association. In case of non-normal data, a Spearman’s correlation is conducted.

#### 2.4.2 Task performance analyses

For paradigm validation, mean accuracy rates (percentage correct trials) and mean response times for correct trials (seconds) from the performance phase were submitted to a one-way repeated measures ANOVA with difficulty level as a repeated measure. Simple effects of difficulty level were assessed using pairwise t-test analyses with a Bonferroni correction. If the assumption of sphericity was not met, a Greenhouse-Geisser correction was applied (Field, 2009).

To confirm that subjective flow scores differ with difficulty level, total flow scores were analyzed using repeated measures ANOVA, with difficulty level as within-subject factor. For both total flow scores as well as component scores, individual ratings were averaged across all trials from each difficulty level. Similarly, we tested whether EC scores were influenced by difficulty level, using the same approach. To assess simple effects of difficulty level on flow and EC scores, pairwise t-test analyses were carried out with a Bonferroni correction. In case of non-sphericity, we have reported Greenhouse Geisser corrected *p*-values.

#### 2.4.3 Anxiety analyses

In order to test whether task performance is influenced by state and/or trait anxiety, we fitted a linear mixed model in which we entered difficulty level, STAI-state scores, STAI-trait scores, and their interactions as fixed effects (accuracy/RT ~ 1+ difficultylevel * STAIstatescore * STAItraitscore + (1 + difficultylevel | subjectnumber). The same procedure was adopted to examine whether state and/or trait anxiety affect people’s subjective engagement score (subjective flow score and effort cost score). Specifically, subjective flow scores were submitted to a mixed model including difficulty level, STAI-state scores, STAI-trait scores, and interactions as fixed effects, while controlling for significant covariates (subjectiveflowscore/ECscore ~ 1+ difficultylevel * STAIstatescore * STAItraitscore + (1 + difficultylevel | subjectnumber).

## 3. Results experiment 1

### 3.1 High correlation between STAI-state and STAI-trait

Spearman’s correlations determined a significant correlation between STAI-state and STAI-trait (ρ(105) = .781, *p* = < .001) (summary statistics can be found in supplementary method section 8.1, table 1).

**Table 1.** State anxiety scores and task performance per condition.

|  | Mean pre | SD pre | Mean post | SD post | Mean difference | SEM difference |
| --- | --- | --- | --- | --- | --- | --- |
| STAI-state score negative | 42.59 | 10.93 | 45.50 | 12.08 | 2.90 | .62 |
| STAI-state score neutral | 39.73 | 11.35 | 40.45 | 10.75 | .72 | .60 |
| STAI-state score positive | 40.48 | 10.53 | 41.19 | 9.50 | .70 | .01 |
| Accuracy rate negative | .66 | .17 | .67 | .19 | .01 | .01 |
| Accuracy rate neutral | .68 | .20 | .71 | .21 | .02 | .01 |
| Accuracy rate positive | .68 | .20 | .71 | .21 | .03 | .21 |
| RT negative | 116.02 | 51.57 | 115.54 | 55.08 | -.48 | 2.43 |
| RT neutral | 129.20 | 86.32 | 130.49 | 76.79 | -1.04 | 4.85 |
| RT positive | 126.84 | 70.63 | 128.55 | 74.13 | 1.71 | 2.45 |

### 3.2 Linear effects of task difficulty on task performance

Average accuracy rate significantly decreased as a function of task difficulty (F(2,56, 271.29) = 292.62, *p* < .001). Specifically, average accuracy rate at each difficulty level was significantly different from all other levels (all *p*s < .001) (Figure 1A). Similarly, correct RT increased significantly as a function of difficulty level (F(2.65, 281.35) = 1182.33, *p* < .001). Post-hoc analyses revealed that RT at each difficulty level was significantly different from that at all other levels (all *p*s < .001) (Figure 1B). These results suggest that participants make more errors and respond more slowly as the task becomes more difficult.

**Figure 1.**
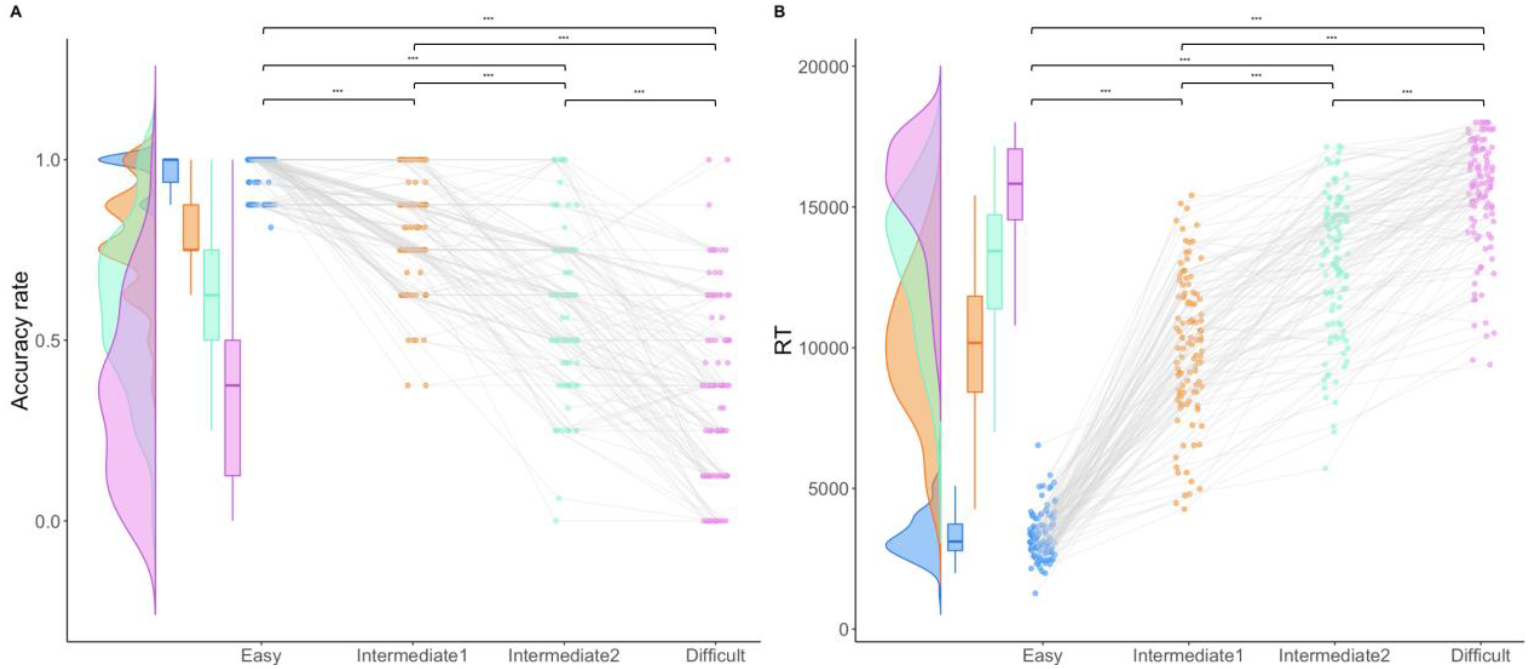
Relationship between difficulty level and task performance. (A) Visualizes the effect of declining accuracy rate as task difficulty increases from easy to difficult levels. (B) Illustrates the increase in RT with increasing task difficulty.

### 3.3 No effect of anxiety on task performance

Due to singularity issues, random slopes for difficulty level were removed from the following two models. First, the mixed-effects regression model confirmed a significant main effect of accuracy rate, with accuracy decreasing as the task became more difficult (β = −.20, SE = .01, t(321) = −24.18, *p <* .*01*). There were no main effects of STAI-state (β <.01, SE < .01, t(107) = .83, *p =* .*41*) or STAI-trait (β < −.01, SE < .01, t(107) = −.10, *p=* .*92*), and there were no interactions between difficulty level and STAI-state (β < .01, SE < .01, t(321) = .89, *p* = .*37*), difficulty level and STAI-trait (β < .01, SE < .01, t(321) = .40, *p* = .*69*), STAI-state and STAI-trait (β < .01, SE < .01, t(107) = .09, *p* = .*93*), or STAI-state, STAI-trait and difficulty level (β < .01, SE < .01, t(321) = .57, *p* = .*57*). Thus, there was no evidence for effects of trait or state anxiety on task accuracy.

Upon fitting the RT model, we encountered the singularity issue. To address this, RT was analyzed using linear models (LM). Parallel with our results in section 3.2, RT significantly increased linearly with increasing difficulty (β = .49, SE = .01, t(1) = 27.57, *p* < .001), such that participants exhibit slower reaction times under more difficult levels. There was no effect of STAI-state (β < .01, SE < .01, t(1) = .25, *p* = .519) or STAI-trait (β < −.01, SE < .01, t(1) = − 2.39, *p* = .066). There were also no significant interaction effects between difficulty level and STAI-state (β < .01, SE < .01, t(1) = .45, *p* = .*712*), difficulty level and STAI-trait (β < −.01, SE < .01, t(1) = −.15, *p* = .*75*), STAI-state and STAI-trait (β < .01, SE < .01, t(1) = .93, *p* = .*355*), or STAI-state, STAI-trait and difficulty level (β < −.01, SE < .01, t(1) = −.30, *p* = .*766*). Additionally, task order was significantly correlated with RT (ρ = .13, *p =* .009) and significantly affected RT (β_Taskorder1_ < −.01, SE_Taskorder1_ = .03, t(3)_Taskorder1_ = −.26; β_Taskorder2_ = .06, SE_Taskorder2_ = .03, t(3)_Taskorder2_ = 2.07; β_Taskorder3_ = −.07, SE_Taskorder3_ = .03, t(3)_Taskorder3_ = −2.68, *p* = .03), reflecting a practice effect. Thus, there was no evidence for effects of state or trait anxiety on reaction times.

### 3.4 Quadratic effect of task difficulty on flow scores and linear effect on effort cost scores

As expected on the basis of previous studies (Csikszentmihalyi, 1990; Ulrich, Johannes, and Grön, 2016), the a priori ANOVA of flow scores confirmed a main effect of task difficulty on the subjective experience of task engagement (F(2.02, 214.09) = 22.74, *p* < .001) (figure 2A). The post-hoc tests revealed a quadratic effect where participants were most engaged at intermediate task levels. Specifically, we observed significantly higher subjective flow scores in the intermediate 1 compared with the easy (*p* < .001) and difficult level (*p* < .001). Furthermore, the intermediate 2 level also showed significantly higher flow scores compared with easy (*p* < .001), and difficult levels (*p* < .001). There was no significant difference between the flow ratings for the intermediate I and intermediate II levels (*p* = .085), and between the easy and difficult levels (*p* = .40).

**Figure 2.**
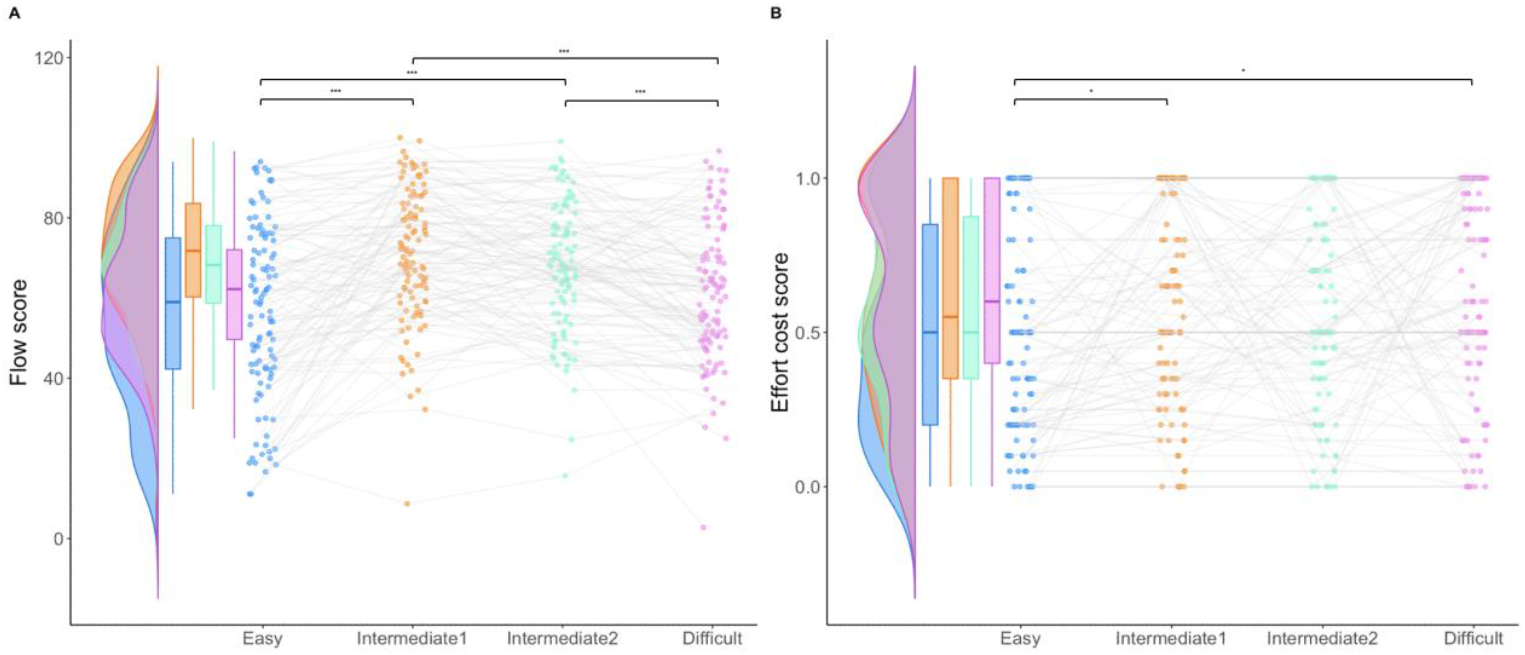
Relationship between difficulty level and effort evaluation. (A) Shows the quadratic effect of subjective flow score over task difficulty. (B) Visualizes the linear increase of effort cost score as task difficulty increases.

There was also a main effect of task difficulty on the effort cost score, which indexes participants’ willingness to avoid performing a difficulty level (F(2.75, 291.21) = 4.8, *p* = .004). This revealed a linear effect, driven by significantly lower EC scores for the easy level compared with both the intermediate 1 level (*p* = .013) and the difficult level (*p* = .012) (figure 2B). No significant differences were observed between the easy and intermediate 2 levels (*p* = .089), intermediate 1 and intermediate 2 levels (*p* = 1.00), intermediate 1 and difficult levels (*p* = 1.000) or intermediate 2 and difficult levels (*p* = 1.00).

These findings demonstrate that subjective flow and effort value ratings vary as a function of task difficulty. In particular, participants were more willing to repeat and thus exert effort for easier tasks than for more difficult (also intermediate) tasks, while also experiencing reduced flow for those easy tasks compared with the more (intermediately) difficult tasks.

### 3.2 Effect of anxiety on subjective flow scores

We hypothesized that state anxiety would be associated with increased effort evaluation and that this effect would be diminished as a function of trait anxiety. To test this, we submitted our two indices of effort evaluation, flow ratings and EC score, to two separate models with state and trait anxiety ratings as predictors.

There was a significant three-way interaction effect of difficulty level, STAI-trait and STAI-state on subjective flow rating (β = −.01, SE = .01, t(107) = −2.52, *p* = .013). There was no main effect of STAI-state scores (β = −.35, SE = .18, t(107) = −1.96, *p* = .053) or STAI-trait scores (β = −.07, SE = .16, t(107 = .44, *p* = .660), and no interaction effects of difficulty level x STAI-trait (β = −.02, SE = .10, t(107) = −.24, *p* = .812), difficulty level x STAI-state (β = .05, SE = .11, t(107) = .45, *p* = .651), or STAI-state x STAI-trait (β = .01, SE = .01, t(107) = .82, *p* = .414). To further understand the three-way interaction, we split the dataset into four subsets based on difficulty level and conducted post-hoc mixed models for each level. Critically, at the easy level only we found a significant interaction effect of STAI-state x STAI-trait (β = .03, SE = .01, t(1) = 2.06, *p* = .042) (figure 3A), whereas neither STAI-state (β = −.42, SE = .30, t(1) = −1.41, *p* = .496) nor STAI-trait (β = .01, SE = .27, t(1) = .04, *p* = .114) showed significant effects. There were no significant effects in the intermediate 1 level: STAI-state (β = −.43, SE = .23, t(1) = .-1.88, *p* = .084), STAI-trait (β = .01, SE = .21, t(1) = .44, *p* = .116), and their (β = .01, SE = .01, t(1) = .68, *p* = .497). At the intermediate 2 level, only the STAI-trait showed a significant main effect (β = −.05, SE = .19, t(1) = .80, *p* = .015), while STAI-state (β = −.36, SE = .21, t(1) = −1.68, *p* = .060) and the interaction (β < −.01, SE = .01, t(1) = −.22, *p* = .826) did not. Finally, in the difficult level there were no significant effects for STAI-state (β = −.28, SE = .24, t(1) = −1.16, *p* = .096), STAI-trait (β = −.02, SE = .21, t(1) = −.11, *p* = .105) or their interaction (β = −.01, SE = .01, t(1) = −1.06, *p* = .294).

**Figure 3.**
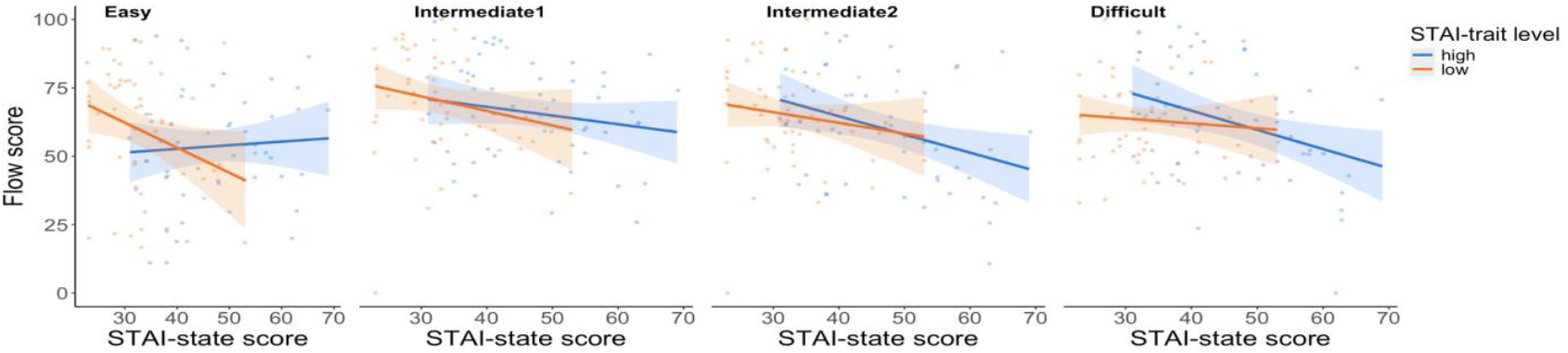
The relationship between subjective flow and anxiety, split by difficulty level. (A) At the easy level, flow score varied depending on the combination of state and state anxiety, where flow decreases state anxiety increases, but only when STAI-trait was low. In contrast, no significant interaction effect of STAI-state and STAI-trait anxiety was observed in the intermediate 1 (B), intermediate 2 (C) nor difficult (D) level.

Effort cost score showed a significant linear effect of difficulty level (β = .04, SE = .01, t(106) = 2.56, *p* = .001) indicating that Effort Cost score increased when the task became more difficult, paralleling the result described in section 3.4. There was no significant effect of STAI-state (β < −.01, SE = .01, t(106) = −.02, *p* = .87) or STAI-trait (β < .01, SE < .01, t(106) = .14, p = .85), nor were there any significant interactions between difficulty level and STAI-state (β < −.01, SE < .01, t(106) = .23, *p* = .986), difficulty level and STAI-trait (β < .01, SE < .01, t(106) = .78, *p* = .555), STAI-state and STAI-trait (β < .01, SE < .01, t(106) = 1.60, *p* = .14), or STAI-state, STAI-trait and difficulty level (β < −.01, SE < .01, t(106) = −.34, *p* = .634) on EC score. This indicates that participant’s willingness to repeat and exert effort was influenced only by task difficulty, and not by anxiety.

### Interim Discussion

While flow induction task performance (accuracy and RTs) scaled linearly with task difficulty, we observed that the effect of task difficulty on subjective flow ratings was inverted U-shaped. Contrary to some (Eysenck et al., 1985) but not all prior literature (Harris & Cumming, 2003), task performance was not modulated by state or trait anxiety.

These results are in line with the previous flow literature (Csikszentmihalyi, 1990; Csikszentmihalyi & Nakamura, 2014; Ulrich et al., 2014) which revealed greater task engagement ratings for intermediately difficult tasks compared with easier and more difficult tasks. Conversely, effort cost scores followed a significant linear trend. This is intriguing because it implies that there is a dissociation between reports of task engagement and measures of task preference.

Importantly, these reports of engagement were modulated by state and trait anxiety, albeit in a different way than we had anticipated. Previous reports (Patzelt et al., 2019) indicated that trait anxiety does not modulate effort avoidance. The current results confirm this finding in showing that trait anxiety does not have a main effect in modulating effort evaluation, while trait anxiety interacted with state anxiety in determining the effect of task difficulty on flow scores, albeit only when the task difficulty was low.

Our original hypothesis was that low trait anxiety coupled with high state anxiety would be associated with increased task engagement as a function of task difficulty. On the contrary, we found that state anxiety was associated with lower flow scores, but only for people with low trait anxiety, and only when the task was easy. This finding aligns with allostatic accounts of stress coping, which propose that chronic stress exposure (associated with high trait anxiety) can dysregulate adaptive responses to acute stressors (state anxiety) (McEwen & Wingfield, 2003). While state anxiety can promote adaptive coping strategies in healthy individuals (Grillon, 2008), trait anxiety has been associated with maladaptive responses and hyper-responsivity to stress (Weger & Sandi, 2018). In this context, individuals with low trait anxiety may have greater cognitive flexibility in deploying adaptive coping mechanisms, allowing them to recognize that low-difficulty tasks are not worth the effort of engagement particularly when state anxiety is high. This aligns with Processing Efficiency Theory (Eysenck & Calvo, 1992), which posits that while trait anxiety reduces processing efficiency, it also motivates increased effort to maintain performance. However, our findings suggest that this compensatory effect may only hold for low-effort tasks. This observation suggests that state anxiety promotes adaptation to low task difficulty, by increasing disengagement (and thus exploration) for the easiest tasks. This would be normative if low task difficulty is associated with high opportunity of exploration, and if state anxiety is a readout of environmental threat that requires more exploration (Agrawal et al 2022). Taken together, these findings suggest that, rather than enhancing engagement under increasing task demands, state anxiety may act as an adaptive disengagement signal in low-effort contexts—particularly for individuals with low trait anxiety who retain cognitive flexibility. This pattern implies that under certain conditions, reduced task engagement in response to heightened state anxiety is not a maladaptive failure of regulation but a strategic shift toward exploratory behavior when cognitive demands are minimal and opportunity costs are high. In this view, anxiety-induced disengagement may serve an adaptive function by reallocating cognitive resources away from low-value tasks toward potentially more informative or rewarding alternatives.

This perspective is also reminiscent of the Yerkes-Dodson law (1908), which suggests that task engagement follows an inverted-U relationship with arousal. While moderate state anxiety may enhance engagement in easy tasks by increasing motivation, excessive arousal during high-effort tasks may disrupt focus and flow. Furthermore, when state anxiety is low, habitual coping strategies in low trait anxiety individuals may function more effectively (Endler, 1997; Endler & Kacovski, 2001), allowing them to maintain engagement regardless of task difficulty. Conversely, when state anxiety is high, the emotional regulation demands increase (Bishop, 2007; Pessoa, 2010), shifting task framing toward threat appraisal and reducing engagement in effortful tasks.

Finally, the effect of anxiety was specific to the flow task and not the effort cost task, emphasizing the dissociation between task engagement, as measured by the flow task and effort cost measures. While the flow induction task relies solely on reports of task engagement, the effort cost task incorporates external rewards (e.g., monetary incentives) to motivate participation in cognitive tasks. The presence of extrinsic rewards may have undermined intrinsic engagement typically associated with task difficulty (Deci, 1971; Cameron & Pierce, 1994). Alternatively, participants may have evaluated the task in relation to task accuracy, which is often tied to the gain of external rewards. This association could naturally have diminished the perceived value of more difficult tasks, where accuracy is lower, reducing the impact of task difficulty on effort cost.

In the next experiment, we move beyond a purely correlative approach by experimentally inducing state anxiety to systematically manipulate its expression across participants. While not a formal intervention assessing pre/post changes, this design allows us to impose controlled variability in state anxiety, thereby enabling us to examine how individual differences in induced state anxiety relate to performance, flow, and effort cost ratings. This approach enhances control over the timing and intensity of anxiety experiences, ensuring consistency across participants and facilitating more precise comparisons between individuals and groups.

## 5. Methods experiment 2

### 5.1 Participants

The sample included 645 English-speaking participants, recruited from the international online portal Prolific. Participated were excluded if they had participated in earlier versions of this experiment. Prior to the experiment, all participants provided an online written informed consent to take part in the study, which was approved by the regional research ethics committee (Commissie Mensgebonden Onderzoek, Arnhem/Nijmegen, CMO2014/288). Each participant performed individually in an online session lasting approximately 90 minutes. All received a monetary reward (€8.66) for their participation with a bonus (€0.00-€1.00) depending on their choices. Participants were removed from the main analyses if they had not completed the study (N = 44), or if they scored 3 SD away of the average accuracy rate and RT in each of the four task levels (N = 29). The final sample consisted of 572 participants (aged 18-77; *M* = 30.90; *SD* = 11.13; 247 women). This study was pre-registered at …

### 5.2 Stimulus Presentation and Data Acquisition

The participants performed the experiment at home at their own computer, following the protocol as in the first experiment. Experimental scripts used to create the task are available on the author’s personal Github page (https://github.com/zceydas/OptEft).

### 5.3 Procedure

#### 5.3.1 Design

The procedure of the second experiment began with the STAI that asked participants about their state and trait anxiety levels (later referred to as STAI-state pre-induction score). Afterwards, participants performed a Sustained Attention to Response Task (SART) (described in section 5.3.3) (later referred to as SART pre-induction score). This was followed by a mood induction (described in section 5.3.2) for which the sample was randomized into three groups: neutral, positive, negative (between-subject factors). These groups correspond to the three mood induction conditions (described in section 5.3.2, mood induction). Specifically, participants were randomly allocated to one of the three conditions in the same ratio, without replacement, with 199 participants in the neutral condition (aged 18-72; *M* = 31.50; *SD* = 11.31; 89 women), 188 participants in the positive condition (aged 18-77; *M* = 31.27; *SD* = 12.96; 87 women), and 185 participants in the negative condition (aged 18-65; *M* = 29.87; *SD* = 9.95; 71 women). Subsequently, participants then completed a second SART and a second STAI-state inventory (later referred as STAI-state and SART post-induction measure). Upon the end of the second STAI-state inventory, the flow induction task started, including flow and effort cost measures to assess effort evaluation of the four difficulty levels (within-subject factors). Additional measures included Need for Cognition (NfC), Positive and Negative Affect Schedule (PANAS, pre- and post-induction measures), and Brief State Rumination Inventory (BSRI, pre- and post-measures). An overview of the design can be seen in figure 4.

**Figure 4.**
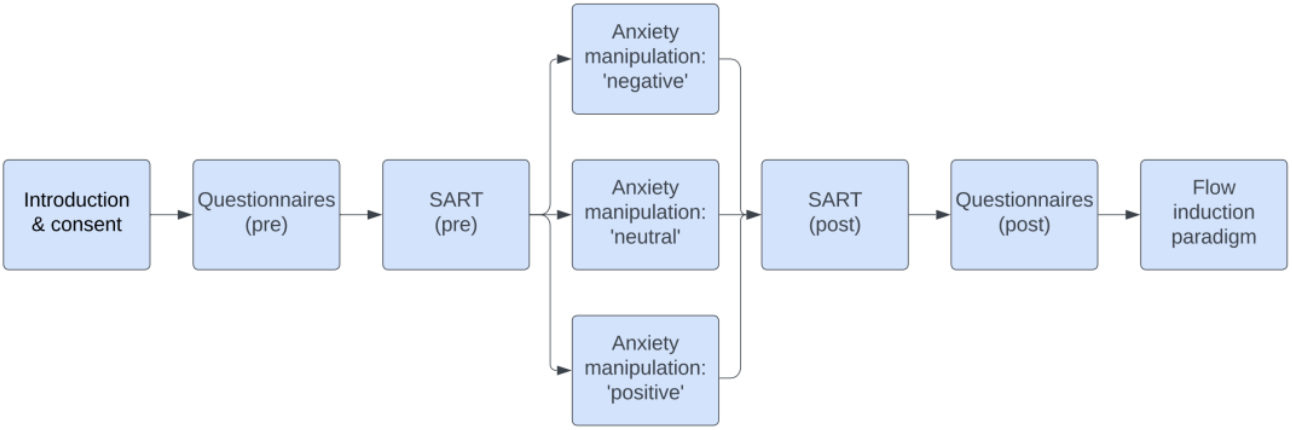
Design of experiment 2.

#### 5.3.2 Mood induction

The mood induction consisted of a free-writing task for 10 minutes about a content depending on the specific group they were allocated to. In the negative group, participants had to recall a negative event that made them feel extremely anxious, had not been resolved yet and was still a source of worry. They had to describe their feelings and give as much detail as necessary to vividly describe the situation. They were instructed to write for the full 10 minutes, and if they did not write a detailed essay, they did not get paid as the experiment builder flagged their participation if the length of the essay was short. After 10 minutes, the participants were automatically directed towards the next phase. The same instructions were given in the positive group, in which participants had to remember, relive and vividly recall an impactful positive event that made them feel extremely happy, and which was still a source of joy for them. The neutral condition consisted of describing an event that took place the day before they participated, which was common to their daily life and was not impactful or emotional. Full instructions can be found in the supplementary method section 9.1.1.

#### 5.3.1. Sustained Attention to Response Task

Participants performed two sustained attention response tasks (SART) both before and after the mood induction to validate that the mood induction was effective, using a non-subjective measure that had previously been shown to be sensitive to state anxiety induction (Aylward & Robinson, 2016). Specifically, in that previous study, participants under threat conditions demonstrated improved accuracy when responding to infrequent distractor stimuli (“no-go” stimuli) and exhibited slower reaction times to frequent target stimuli (“go” stimuli). These changes suggested that anxiety enhances motor response inhibition, a mechanism that might be overactive in individuals with clinical anxiety. The findings indicate that anxiety reliably impacts both accuracy and reaction time, suggesting the SART’s potential as a non-subjective measure of anxious responding with clinical relevance.

Participants had to respond to the target stimuli (go trials) and withhold responses to (infrequent) non-target stimuli (no-go trials) in this task. Go and no-go cues were presented on the screen in a subsequent, random order. When a go cue was presented, participants needed to press the spacebar as soon as possible, whereas for the no-go cue they had to withhold their behavioral response of pressing the spacebar. In this experiment, participants started with ten practice trials of eight go and two no-go cues. Subsequently, participants performed four blocks each consisting of 47 go trials and 5 no-go trials. Reaction time (RT) and accuracy rate were measured for both go trials (correct responses) and no-go trials (incorrect responses). Outliers (RT > 1000 ms) were rejected (Righi, Mecacci, & Viggiano, 2009).

### 5.4 Statistical analyses

We employed the same analysis pipeline as in experiment 1, which included inventory, behavioral, and inventory-behavioral analyses. However, in all models STAI-state post scores were used as predictor. Additionally, for the current experiment we incorporated additional analyses to validate the mood induction. Specifically, we conducted linear mixed models predicting STAI-state score, and accuracy and RT of the SART by time (pre and post), mood induction group (negative, positive, neutral), and their interactions (STAI-state score/accuracy/RT ~ 1 + time * condition + (1 | subjectnumber). There was no multicollinearity between predictors in this dataset (see supplementary section 9.2, table 7-8 for an overview of the VIF scores). The results of the inventory analyses and behavioral analyses can be found in supplementary section 9.2.3 and 9.2.4. VIF scores and (relevant) model estimates of all four flow subcomponents (involvement, liking, ability, time) as well as models using condition instead of STAI-state post scores can be found in supplementary section 9.1, table 9-10, 9.2, table 11-16).

## 6. Results section experiment 2

### 6.2 Negative mood induction increased state anxiety

The efficacy of the mood induction was demonstrated by showing a significant interaction effect of time (pre and post mood induction) and condition (β_preneutral_ = .73, SE_preneutral_ = .24, t(572)_preneutral_ = 3.04; β_prepositivel_ = −.36, SE_prepositive_ = .24, t(572)_prepositive_ = −1.53, *p* < .01). This effect was driven by greater increases in the STAI-state score from pre-to post-induction in the negative mood condition (β = 1.45, SE = .31, t(185) = 4.66, p < .001), but not in the neutral (β = .36, SE = .30, t(199) = 1.20, p = .231) or positive mood condition (β = .35, SE = .26, t(188) = 1.34, p = .182; figure 7A and 7B; table 1). In addition, there were significant main effects of time (β = .72, SE = .17, t(572) = 4.27, *p* < .001) and condition (β_neutral_ = 2.39, SE_neutral_ = .17, t(572)_neutral_ = 3.97; β_positive_ = −1.57, SE_positive_ = .59, t(572)_positive_ = −2.65, *p* < .001) on STAI-state score.

SART accuracy rate did not show a significant interaction effect (β_preneutral_ = −.01, SE_preneutral_ = .004, t(563)_preneutral_ = −1.20; β_prepositivel_ = .002, SE_prepositive_ = .005, t(563)_prepositive_ = .44, *p* = .481). There was a significant effect of time (β = .69, SE = .003, t(563) = 3.05, p = .002), increasing from pre-to post-mood-induction (Mpre = .68, SDpre = .19; Mpost = .70, SDpost = .21), perhaps reflecting a practice effect (Aylward and Robinson, 2017), see also section 6.2). There was no main effect of condition (β_neutral_ = −.02, SE_neutral_ = .01, t(563)_peutral_ = −1.79; β_positivel_ = 01, SE_positive_ = .01, t(563)_positive_ = .60, *p* =.193; figure 7 C, D).

Reaction time (RT) of the SART go trials was not affected by condition or time. We did not find any significant effects (main effect of time: β = .12, SE = 1.02, t(554) = .12, p = .91; main effect of condition: β_neutral_ = −.9.00, SE_neutral_ = 4.01, t(555)_neutral_ = −2.23; β_positive_ = 6.00, SE_positive_ = .3.94, t(555)_positive_ = 1.52, *p* = .074; time x condition interaction: β_preneutral_ = −.36, SE_preneutral_ = 1.46, t(554)_preneutral_ = −.25; β_prepositivel_ = −.37, SE_prepositive_ = 1.43, t(554)_prepositive_ = −.26, *p* = .88) (figure 7 E, F). Thus, there was no evidence for an effect of negative mood induction on SART performance.

There was a significant increase in STAI-state scores after the negative mood induction only, STAI-state scores did not change for the neutral and positive condition. The groups did not differ in their accuracy rate; participants had increased accuracy rates from pre to post mood-induction overall. RT did not differ per group or time.

### 6.2 No effect of anxiety on task performance

Random slopes for difficulty level were removed from the following models because of singularity. As seen previously (Exp 1), accuracy rates decreased significantly with task difficulty (β = −.21, SE < .01, t(1716) = −59.83, *p <* .*001*). Also, there were no significant main effects of STAI-state (β < .01, SE < .01, t(571) = .1.19, *p* = .236) and STAI-trait (β < .01, SE < .01, t(571) = −.38, *p* = .707). Furthermore, there were no interaction effects between task difficulty and STAI-state (β < .01, SE < .01, t(1713) = 1.65, *p* = .099), task difficulty and STAI-trait (β < .01, SE < .01, t(1713) = −.37, *p* = .709), STAI-state and STAI-trait (β < .01, SE < .01, t(571) = .44, *p* = .658), or STAI-state, STAI-trait and difficulty level (β < .01, SE < .01, t(1713) = −1.60, *p =* .*109*).

RT increased linearly with increasing task difficulty (β = .44, SE = .01, t(539) = 60.93, *p* < .001). There were no main effects of STAI-state (β < −.01, SE < .01, t(1) = −.36, *p* = .730) or STAI-trait (β < −.01, SE < .01, t(1) = 2.62, *p* = .106), and there were no interactions between difficulty level and STAI-state (β < −.01, SE < .01, t(1) = −.30, *p* = .763), difficulty level and STAI-trait (β < −.01, SE < .01, t(1) = −.63, *p* = .305), STAI-state and STAI-trait (β < .01, SE < .01, t(1) = .125, *p* = .927), or STAI-state, STAI-trait and difficulty level (β < .01, SE < .01, t(1) = .37, *p* = .372). Thus, as was the case for experiment 1, task performance did not vary with state- or trait anxiety.

### 6.3 Effect of mood induction on effort evaluation

STAI-state anxiety was significantly associated with lower flow scores, regardless of the task difficulty (β = −.27, SE = .07, t(571) = −4.14, *p* < .001) (figure 5A). In addition, there was a significant interaction effect of STAI-state and STAI-trait (β = .01, SE < .01, t(571) = 2.61, *p* = .009). To explore this effect, we split the dataset into two subsets based on the median score of STAI-state and conducted post-hoc mixed models for each subset. In the low STAI-trait subset we found a significant effect of STAI-trait (β = −.43, SE = .08, t(299) = −5.10, *p* < .001), due to higher state anxiety being associated with lower flow scores. This effect of state anxiety on flow was present but less steep in the high STAI-trait subset (β = −.17, SE = .08, t(273) = − 2.22, *p* = .033) (figure 5B). There was no significant effect of task difficulty (β = .56, SE = .48, t(571) = 1.18, *p* = .240), STAI-trait (β = −.08, SE = .07, t(571) = −1.47, *p* = .144), or significant interactions between task difficulty and STAI-state (β = −.03, SE = .06, t(571) = −.52, *p* = .602), task difficulty and STAI-trait (β = −.05, SE = .04, t(571) = −1.26, *p* = .208), and task difficulty, STAI-state and STAI-strait (β < −.01, SE < .01, t(571) = −.06, *p* = .954).

**Figure 5.**
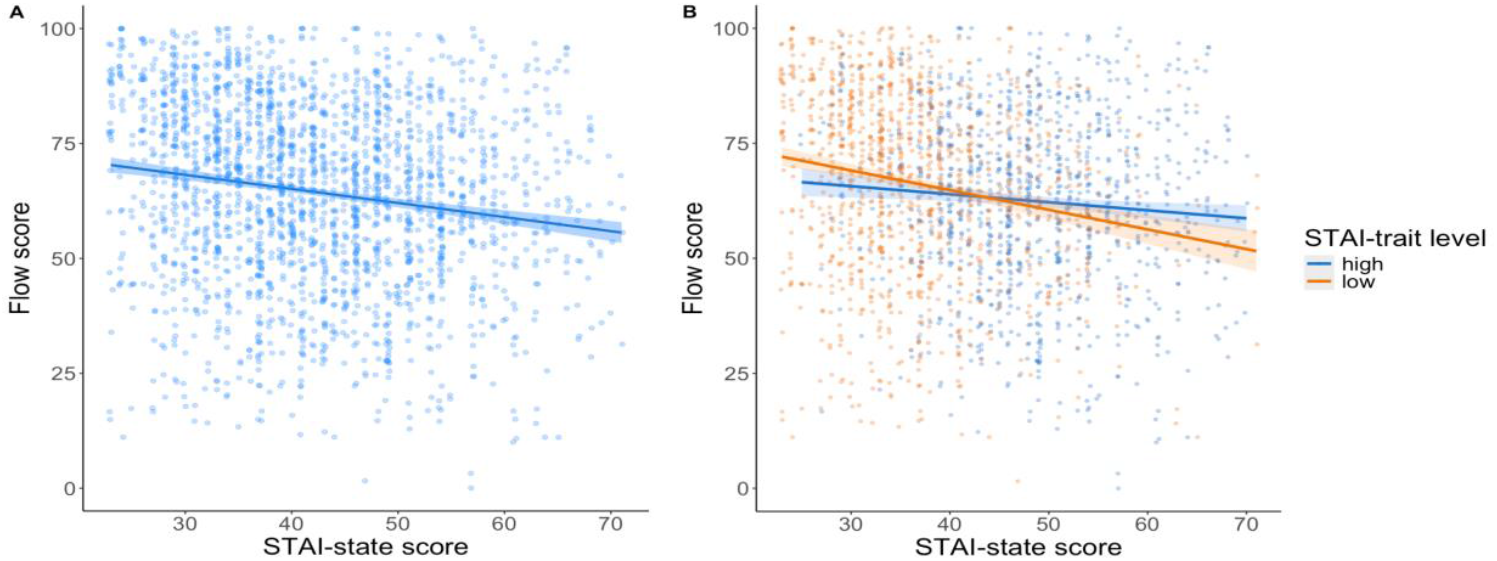
The effect of anxiety on flow score (A) This figure visualizes a negative relationship between flow score and STAI-state score. (B) A higher STAI-state results in lower flow scores only when STAI-trait levels are low. This relationship is less strong when STAI-trait levels are high.

As in experiment 1, effort cost varied with task difficulty (β = .03, SE = .01, t(571) = 5.25, p < .001). Critically, there was also a significant main effect of STAI-state (β < .01, SE < .01, t(571) = 2.28, *p* = .012; figure 6A) and STAI-trait (β < −.01, SE < .01, t(571) = −2.52, *p* < .023; figure 6B). In line with the findings of experiment 1, there were no significant interactions between task difficulty and STAI-state (β < .01, SE = .01, t(571) = −1.04, *p* = .299), task difficulty and STAI-trait (β < .01, SE < .01, t(571) = .65, *p* = .515), STAI-state and STAI-trait (β < −.01, SE < .01, t(571) = −.14, *p* = .890) and no three-way interaction between task difficulty, condition, and STAI-trait scores (β < −.01, SE < .01, t(571) = −.04, *p* = .970).

**Figure 6.**
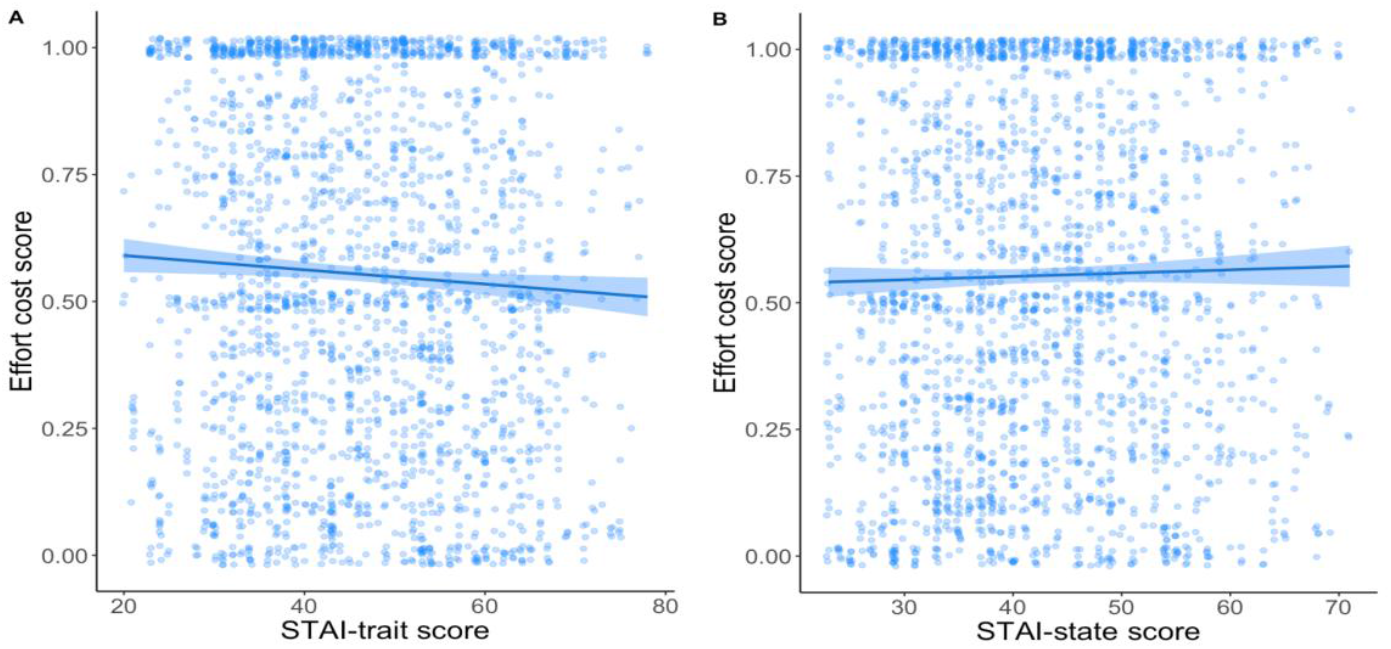
The relationship between effort cost scores and anxiety This figure illustrates opposing effects of trait and state anxiety: (A) effort costs decrease as STAI-trait scores increase, while (B) effort costs increase with higher STAI-state scores.

Overall, there was less willingness to repeat difficult tasks. However, this willingness interacted with both state and trait anxiety, so that state anxiety was associated with lower flow to a greater degree in people with lower trait anxiety.

### Interim Discussion

In this experiment, we have taken a comprehensive approach to understand the interplay between anxiety, cognitive performance, and task engagement. Diverging from prior experiments, we included a mood induction phase to induce variability in state anxiety.

The main effects of state anxiety on flow scores and effort cost scores indicate that higher state anxiety is associated with reduced flow and higher effort cost. As was the case in experiment 1, the effect of state anxiety on flow scores depended on trait anxiety, so that it was reduced in those with higher trait anxiety. Unlike the effect in experiment 1, this effect in experiment 2 was not restricted to the easiest task levels, but was present across all levels of task difficulty. The absence of a significant interaction effect between state, trait anxiety and task difficulty on flow scores in this experiment could suggest that the specific mood induction used here may have buffered or overshadowed the more nuanced effects of anxiety on task performance.

However, it should be noted that participants in Experiment 2 had overall higher baseline trait anxiety levels compared with participants in Experiment 1 (supplementary section 9.2.1). Importantly, the absence of a task difficulty effect in this context is unlikely to reflect a stronger trait effect. Rather, it is more plausibly explained by a dominant state anxiety effect that is suppressed under high trait anxiety. After all, the interaction pattern reflects an abolition of the state anxiety effect in individuals with higher trait anxiety, rather than a straightforward amplification of trait-driven effects.

In terms of effort cost, all experiments consistently demonstrated that participants were less inclined to exert effort as task difficulty increased. However, while participants with high state anxiety showed a decreased willingness to repeat tasks, reflecting an aversion to effortful tasks under heightened situational stress, individuals with high trait anxiety demonstrated an increased willingness to repeat tasks, potentially driven by compensatory mechanisms or a greater tolerance for sustained engagement, even in the face of task difficulty.

## General Discussion

The present series of experiments uncovered a systematic relationship between trait anxiety, state anxiety, and task engagement across two independent experiments. As task difficulty increased, participants were consistently less willing to exert effort, aligning with demand avoidance patterns (Kool, McGuire, Rosen, & Botvinick, 2010). The quadratic effect of task difficulty on flow ratings was replicated across both experiments, confirming the inverted U-shaped relationship proposed by Csikszentmihalyi (1990), in which maximum engagement occurs at an optimal level of challenge. These findings underscore a dissociation between effort cost evaluations and subjective engagement: while participants were most engaged by intermediately difficult tasks, their willingness to repeat tasks declined linearly with difficulty.

Our initial hypothesis stated that state anxiety would enhance task engagement with increasing task difficulty, but that this effect would be diminished in individuals with high trait anxiety due to impaired deployment of adaptive coping mechanisms. However, results from both experiments contradicted this expectation. In Experiment 1, state anxiety was associated with *reduced* flow, but only in individuals with low trait anxiety, and only when the task was easy. There was no evidence that state anxiety increased engagement at higher difficulty levels. State anxietymotivated disengagement under conditions of low cognitive demand, and this effect was attenuated by trait anxiety. This result aligns with allostatic accounts of stress coping (McEwen & Wingfield, 2003), which propose that chronic stress exposure (indexed by high trait anxiety) impairs adaptive responses to acute stress (state anxiety). While Processing Efficiency Theory (Eysenck & Calvo, 1992) suggests that anxiety may motivate compensatory effort, our results indicate that this compensatory mechanism may be limited to specific contexts—particularly under low task difficulty where disengagement is normatively adaptive. This is consistent with the idea that state anxiety may increase the perceived value of disengagement in unchallenging contexts (Agrawal et al., 2022), potentially signaling the need to shift from exploitation to exploration.

Experiment 2 introduced a mood induction procedure to systematically manipulate state anxiety. As in Experiment 1, higher state anxiety was associated with reduced flow. This effect again depended on trait anxiety: individuals with high trait anxiety were less sensitive to changes in state anxiety, showing a general dampening of the state anxiety effect. Importantly, this interaction was present across all levels of task difficulty, rather than being restricted to easy tasks. Notably, participants in Experiment 2 had higher baseline trait anxiety than those in Experiment 1, which may have contributed to a stronger trait-driven suppression of the state anxiety effect. The result was a clearer pattern in which low state and low trait anxiety jointly predicted the highest engagement. These findings highlight that while state anxiety may negatively influence engagement, its effect is contextually gated by trait anxiety.

Together, the two experiments reveal that state anxiety is not reliably associated with increased engagement, even under high cognitive demand. Instead, high trait anxiety appears to attenuate the effects of acute state anxiety, thus disrupting task disengagement.

Across both experiments, the dissociation between flow and effort cost was consistently observed. While flow ratings were modulated by anxiety and task difficulty, effort cost scores were predominantly driven by task difficulty alone. Participants became less willing to exert effort as task demands increased, regardless of anxiety level. This is consistent with the idea that effort cost is shaped more by extrinsic motivational factors—such as reward or accuracy— than by internal engagement states (Deci, 1971; Cameron & Pierce, 1994; Dunn et al., 2019). In difficult tasks, where accuracy tends to be lower, participants may devalue the task more quickly, thereby reducing willingness to engage regardless of intrinsic flow experiences.

An important methodological feature of Experiment 2 was the inclusion of a mood induction to experimentally manipulate state anxiety. This procedure successfully introduced controlled variability in state anxiety, but may have also buffered or overshadowed more subtle interactions with task difficulty seen in Experiment 1. The lack of significant interaction between anxiety and performance measures in Experiment 2 may reflect this dampening. Differences from prior findings using stronger physiological stressors (e.g., electric shocks; Aylward & Robinson, 2016) may also account for the relatively moderate effects observed here, as our autobiographical essay method likely elicited a less acute stress response.

Finally, we note a discrepancy between the current findings and those of Sayali, Heling, & Cools (2023), who reported a linear increase in flow with task difficulty. This divergence likely reflects methodological differences, specifically trial number per condition. Sayali et al. used two blocks per difficulty level, allowing for greater performance improvement over time, especially in high-difficulty conditions. This performance gain may have enhanced engagement and shifted the flow–difficulty relationship from an inverted-U to a linear trend. In the current study, each difficulty level was only tested once, minimizing the opportunity for performance-related gains to modulate engagement.

