## Supplemental for "Anxiety state-related task disengagement varies with trait anxiety"

### 8.1 Supplementary method section experiment 1

*8.1.1. Additional psychological inventories*

The second inventory included was the Need for Cognition (NfC), to assess the effect of pre-existing differences in motivation to invest cognitive effort in our model (Cohen, Stotland, & Wolfe, 1955). The third inventory executed was the Positive Affect and Negative Affect Scale (PANAS), developed by Watson, Clark and Tellegen (1988), which contains two subscales: positive and negative affect. Each subscale consists of ten emotional words that must be rated on a scale from 1 to 5 (1 being “not at all”, and 5 being “very much”), used to measure the current mood of the participants. Lastly, participants completed the Brief State Rumination Inventory (BSRI), developed by Marchetti et al. (2018). This questionnaire measures state rumination and was used to see ruminative behavior could affect flow experiences and effort cost behavior. It contains eight statements in which participants had to declare how much each statement applied to them by reporting on a 100-mm visual analog scale ranging from 0 to 100 (0 being “completely disagree”, and 100 being “completely agree”). In addition, the relationship between all inventories was explored by means of a pearson’s correlation (see supplementary section 8.2, figure 2, for all scatterplots and figure 3 for a heatmap).

8.1.2. *Instructions effort cost*

“In the next screen, we will ask you to indicate the MINIMUM amount of money you would be willing to earn in exchange for REPEATING the type of questions you just experienced. After you indicate the minimum amount, you are willing to earn for repeating a type of question, we will randomly draw a number between 0 and 1 euro. If the number we draw is greater than yours, then you will REPEAT a type of question in exchange for the amount you drew. If the number we draw is equal to or less than yours, you will NOT REPEAT a question type and earn nothing. It’s actually really simple: would you like to repeat a question type? Make a smaller bid. You don’t want to repeat a question type? Make a larger bid. Please note that your ACCURACY in solving the arithmetic questions DOES NOT AFFECT whether you will earn the extra amount as long as you TRY YOUR BEST and continue solving each block in your mind and without guessing or using any other tools.”.

| | STAI-state | STAI-trait | BSRI | PANAS negative | PANAS positive | NfC |
|---|---|---|---|---|---|---|
| Mean | 39.93 | 43.86 | 313.72 | 21.24 | 31.47 | 57 |
| SD | 11.43 | 12.20 | 179.96 | 7.94 | 7.44 | 11.59 |
| Median | 38 | 44 | 291 | 20 | 32 | 58 |

Table 1. Summary statistics of inventories.

| Variables | Difficulty level | STAI-state | STAI-trait | STAI-state x STA-Trait | Difficulty level x STAI-state | Difficulty level x STAI-trait | Difficulty level x STAI-state x STAI-trait | Covariate task order | Covariate age |
|---|---|---|---|---|---|---|---|---|---|
| RT | 1.57 | 2.68 | 2.48 | 1.17 | 2.72 | 2.50 | 1.73 | 1.05 | |
| Accuracy rate | 1.55 | 2.68 | 2.48 | 1.17 | 2.68 | 2.48 | 1.72 | | |
| Flow score | 1.55 | 2.71 | 1.60 | 1.18 | 2.69 | 2.49 | 1.72 | | 1.19 |
| EC score | 1.55 | 1.71 | 1.50 | 1.71 | 1.71 | 1.50 | 1.73 | | |

Table 2. VIF scores of all models.

| Variables | STAI-state | STAI-trait | STAI-state x STA-Trait |
|---|---|---|---|
| Easy subset | 2.68 | 2.48 | 1.17 |
| Intermediate 2 subset | 2.68 | 2.48 | 1.17 |
| Intermediate 2 subset | 2.68 | 2.48 | 1.17 |
| Difficult subset | 2.68 | 2.48 | 1.17 |

Table 3. VIF scores of the difficulty sub models

| Variables | STAI-state x STA-Trait | Difficulty level x STAI-state x STAI-trait |
|---|---|---|
| Involvement | 1.38 | 1.86 |
| Liking | 1.18 | 1.72 |
| Ability | 1.34 | 1.89 |
| Time | 1.19 | 1.73 |

Table 4. VIF scores of all subjective flow subcomponent models: relevant interaction effects

### 8.2 Supplementary result section experiment 1

#### *8.2.1. Additional psychological inventories*

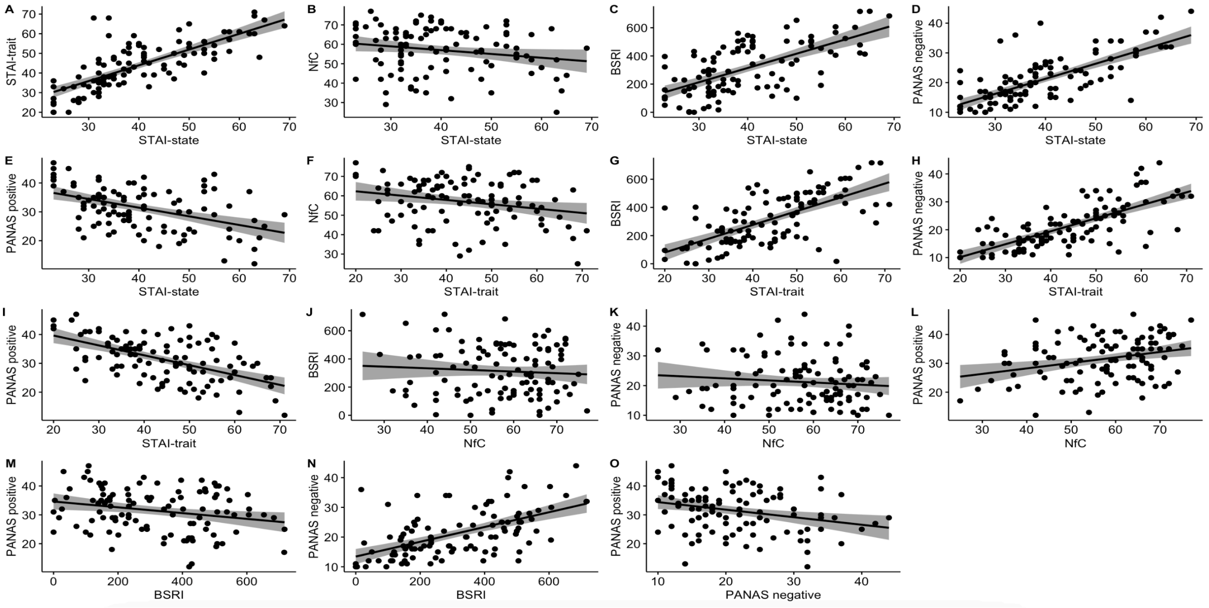


Figure 1. Scatterplots of relationship between the inventories. Each dot represents a single participant.

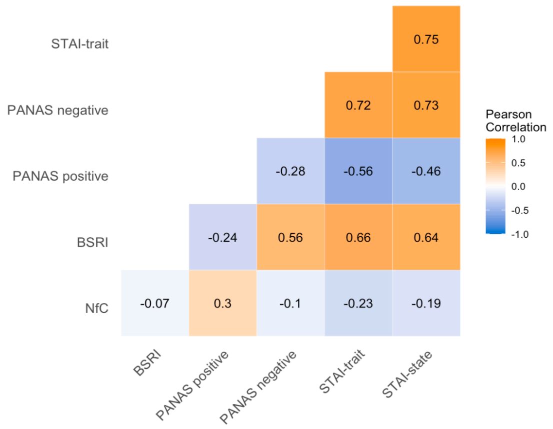


Figure 2. Heatmap showing the correlation coefficients per inventory. Correlation is significant at the .05 level.

#### *8.2.2. Effect of anxiety on SART RT but not on accuracy rate*

To explore a relationship between state anxiety and performance on the SART, we performed two pearson's correlations between STAI-state score and accuracy rate (no-go trials) as well as

RT (go-trials). There was a significant negative correlation between STAI-state and RT (r(103) = -.19, *p* = *.049*), which was not found for accuracy rate (r(103) = -.11, *p* = *.263*). So, in this experiment higher levels of state anxiety decrease response speed but did not affect response inhibition see figure 3.

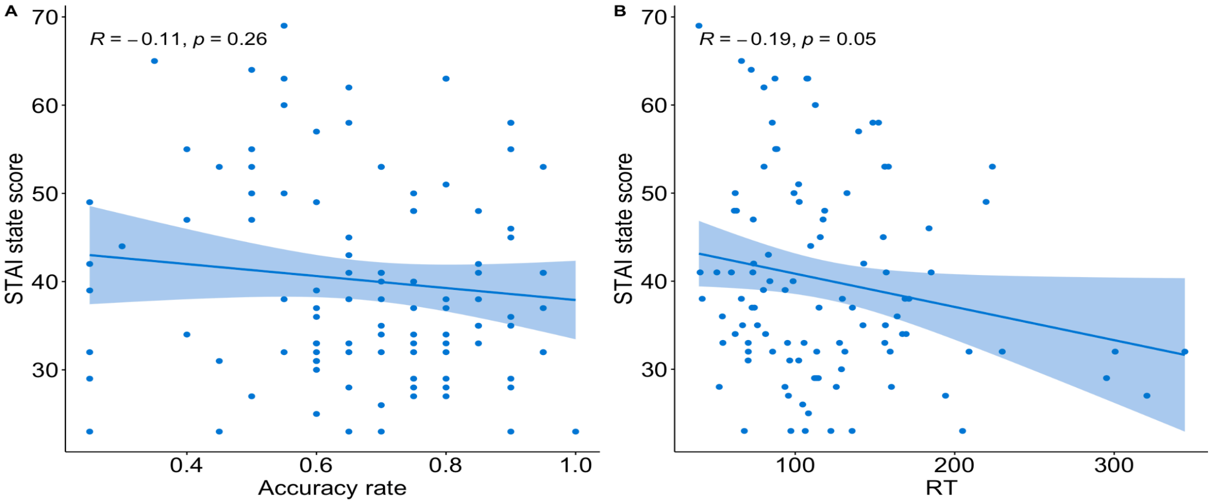


Figure 3. Scatterplots visualizing the relationship between STAI-state and task performance. (A) a negative correlation between STAI-state score and SART accuracy rate. (B) a negative correlation between STAI-state score and SART RT. Each dot represents a single participant.

| Predictors | Estimate | SD | t value | P value | Df |
|---|---|---|---|---|---|
| Involvement: STAI-state x STAI-trait | .03 | .02 | 1.48 | .14 | 106 |
| Involvement: Difficulty level x STAI-state x STAI-trait | -.01 | .01 | -1.41 | .16 | 106 |
| Liking: STAI-state x STAI-trait | .002 | .01 | .14 | .89 | 107 |
| Liking: Difficulty level x STAI-state x STAI-trait | -.01 | .01 | -2.15 | .034 | 107 |
| Ability: STAI-state x STAI-trait | -.01 | .01 | -1.16 | .250 | 107 |
| Ability: Difficulty level x STAI-state x STAI-trait | -.01 | .01 | -1.44 | .153 | 107 |
| Time: STAI-state x STAI-trait | .004 | .01 | .31 | .76 | 1 |
| Time: Difficulty level x STAI-state x STAI-trait | -.02 | .01 | -1.64 | .104 | 107 |

Table 5. Flow subcomponent model output: relevant interaction effects

| Predictors | Estimate | SD | t value | P value | Df |
|---|---|---|---|---|---|
| Easy: STAI-state | -.48 | .37 | -1.28 | .381 | 1 |
| Easy: STAI-trait | .26 | .34 | .77 | .749 | 1 |
| Easy: STAI-state x STAI-trait | .02 | .02 | 1.24 | .217 | 1 |
| Intermediate 1: STAI-state | -.27 | .33 | -.81 | .435 | 1 |
| Intermediate 1: STAI-trait | .07 | .29 | -.23 | .18 | 1 |
| Intermediate 1: STAI-state x STAI-trait | < .01 | .02 | .23 | .153 | 1 |
| Intermediate 2: STAI-state | -.37 | .32 | -1.16 | .192 | 1 |
| Intermediate 2: STAI-trait | -.18 | .29 | -.61 | .018 | 1 |
| Intermediate 2: STAI-state x STAI-trait | < -.01 | .02 | -.15 | .885 | 1 |
| Difficult: STAI-state | -.15 | .32 | -.47 | .271 | 1 |
| Difficult: STAI-trait | -.21 | .29 | -.73 | .076 | 1 |
| Difficult: STAI-state x STAI-trait | -.02 | .02 | -1.47 | .145 | 1 |

Table 6. Flow subcomponent liking model output: three-way interaction split

### 9.1 Supplementary method section experiment 2

#### *9.1.1. Introduction text of anxiety induction*

(Instructions positive group: *"Please remember, relive, and vividly recall a positive event that makes you feel extremely happy. Choose an event that has been impactful and is still a source of joy for you. Please give as much details as necessary to vividly describe the situation and try to describe your feelings. You will have 10 minutes to complete this task. You must write for the full 10 minutes. The study will automatically continue when the 10 minutes is over. Note that if you haven't written a detailed essay for this section, your participation will be flagged and you may not get paid"*. Instructions negative group: *"Please remember, relive, and vividly recall a negative event that makes you feel extremely anxious. Choose an event that has not been resolved and is still a source of worry for you. Please give as much details as necessary to vividly describe the situation and try to describe your feelings. You will have 10 minutes to complete this task. You must write for the full 10 minutes. The study will automatically continue when the 10 minutes is over. Note that if you haven't written a detailed essay for this section, your participation will be flagged, and you may not get paid"*. Instructions neutral group: *"Please remember, relive, and vividly recall one of the events that took place yesterday. Choose an event that is common to your daily life and is not impactful or emotional to you. Please give as much details as necessary to vividly describe the situation and try to describe your feelings. You will have 10 minutes to complete this task. You must write for the full 10 minutes. The study* *will automatically continue when the 10 minutes is over. Note that if you haven't written a detailed essay for this section, your participation will be flagged, and you may not get paid".*)

*9.1.2. Baseline STAI score analysis*

To test whether there were baseline differences between the STAI-state inventory conducted in experiment 1 and the STAI-state pre anxiety induction inventory (full dataset; all conditions), we performed an independent t-test. The same test was run to test for potential differences in STAI-trait scores.

*9.1.3. Anxiety induction correlation analyses*

We examined whether state anxiety would affect performance of the SART. To this end, we performed a Pearson correlation on accuracy rate of the no-go trials with STAI-state score and RT of the go trials with STAI-state score, for the pre as well as post anxiety induction measures.

*9.1.4. Inventory-behavioral & motivational-anxiety models including STAI-state score*

To further explore the effect of anxiety on task performance as well as subjective engagement score and effort cost, the following models include STAI-state post score (accuracy/RT/subjectiveflowscore/ECscore ~ 1+ taskdifficultylevel * Condition * STAItraitscore + NfCscore + taskorder + age + yearsofeducation + gender + (1 + taskdifficulty level | subjectnumber).

| Variable | Time | Condition | TimeCondition |
|---|---|---|---|
| STAI-state score | 1.00 | 1.00 | 1.00 |
| Accuracy rate | 1.00 | 1.00 | 1.00 |
| RT | 1.00 | 1.00 | 1.00 |

Table 7. VIF scores of models: mood induction

| Variables | Difficulty level | Condition | STAI-trait | Condition x STAI-trait | Difficulty level x condition | Difficulty level x STAI-trait | Difficulty level x condition x STAI-trait | Covariate years of education | Covariate age | Covariate gender |
|---|---|---|---|---|---|---|---|---|---|---|
| Accuracy rate | 1.40 | 1.80 | 1.79 | 1.02 | 1.80 | 1.79 | 1.41 | 1.00 | | |
| RT | 1.39 | 1.79 | 1.78 | 1,02 | 1.77 | 1.76 | 1.40 | | | |
| Flow scores | 1.39 | 1.84 | 1.83 | 1.02 | 1.81 | 1.79 | 1.42 | | 1.08 | 1.02 |
| EC scores | 1.40 | 1.83 | 1.80 | 1.02 | 1.80 | 1.79 | 1.41 | | 1.07 | |

Table 8. VIF scores of models: STAI-state post score

| Variables | STAI-state x STAI-trait | Difficulty level x STAI-state x STAI-trait |
|---|---|---|
| Involvement | 1.10 | 1.50 |
| Liking | 1.03 | 1.43 |
| Ability | 1.38 | 1.78 |
| Time | 1.02 | 1.41 |

Table 9. VIF scores of all subjective flow subcomponent models: relevant interaction effects

| Variables | Difficulty level | Condition | STAI-trait | Condition x STAI-trait | Difficulty level x condition | Difficulty level x STAI-trait | Difficulty level x condition x STAI-trait | Covariate age | Covariate years of education | Covariate gender |
|---|---|---|---|---|---|---|---|---|---|---|
| Accuracy rate | 1.00 | 1.01 | 1.01 | 1.01 | 1.01 | 1.01 | 1.01 | | 1.01 | |
| RT | 1.00 | 1.01 | 1.01 | 1.01 | 1.01 | 1.01 | 1.01 | | | |
| Flow score | 1.00 | 1.03 | 1.07 | 1.02 | 1.02 | 1.01 | 1.02 | 1.07 | | 1.04 |
| EC score | 1.00 | 1.01 | 1.06 | 1.01 | 1.01 | 1.08 | 1.01 | 1.06 | | |

Table 10. VIF scores of models: condition

### 9.2 Supplementary result section experiment 2

*9.2.1 No significant baseline differences in STAI-state but STAI-trait score is significantly higher in the dataset of experiment 2*

The t-tests did not show a significant difference in baseline STAI-state score (pre score for the second dataset) ($M_{first}$ = 40.64, $SD_{first}$ = 11.55; $M_{second}$ = 40.91, $SD_{second}$ = 11.37; t(665.21) = .44, *p* = .661*)*. However, there was a significant difference between STAI-trait score ($M_{first}$ = 44.65, $SD_{first}$ = 12.32; $M_{second}$ = 46.37, $SD_{second}$ = 12.61; t(682.07) = 2.74, *p* = .006*)*. Thus, we ruled out that participants score significantly different for STAI-state in the first dataset compared to the second, but participants in the second experiment have a significantly higher STAI-trait score.

*9.2.2. Negative correlation between STAI-state score and SART acc but not for RT*

We found a significant negative correlation between accuracy rate (no-go trials) and STAI-state before the anxiety induction (r = -.14, p = .001; Figure 4A) as well as after the anxiety induction (r = -.20, p < .001; Figure 4B). This indicates that higher STAI-state scores were associated with lower accuracy rates on SART across all participants. Additionally, there was no significant correlation between RT (go trials) and STAI-state scores before the anxiety induction (r = -.04, p = .318; Figure 4C) or after (r = -.05, p = .267) (Figure 4D). This demonstrates that state anxiety impairs accuracy in SART with no influence on response speed.

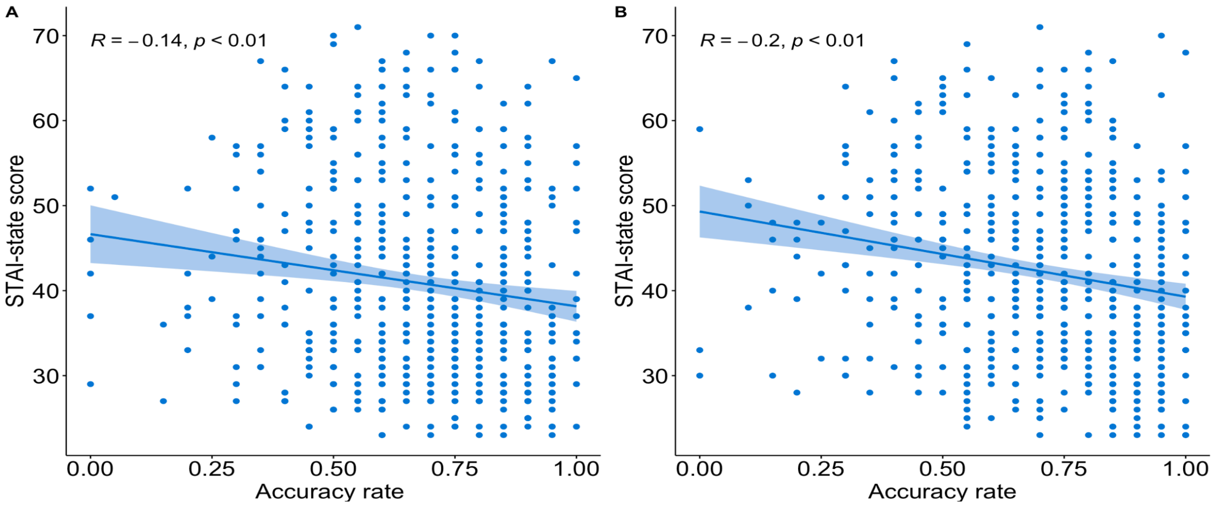

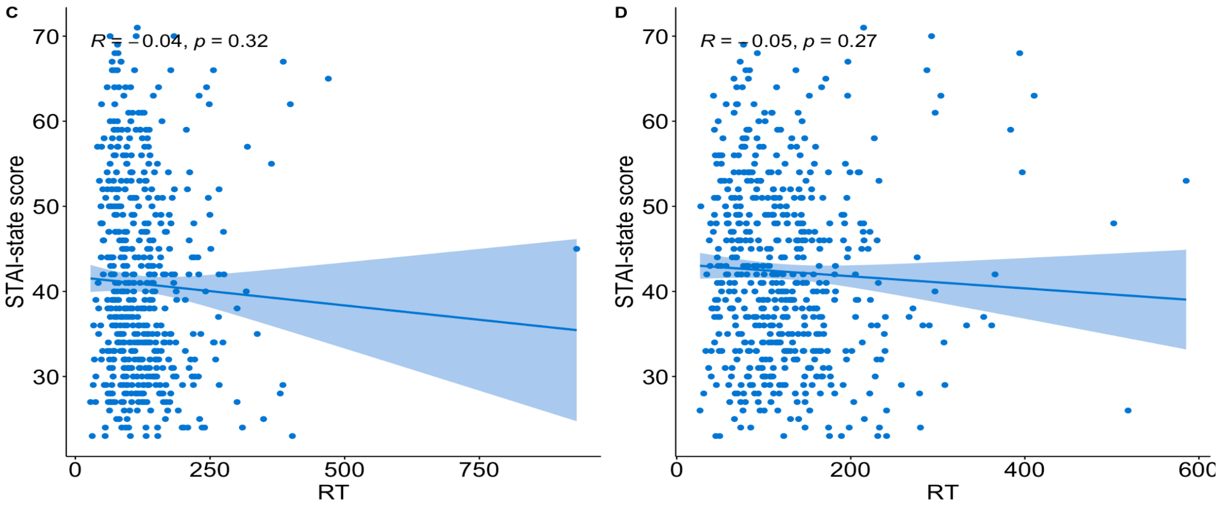


Figure 4. Scatterplots showing the relationship between STAI-state score and task performance. Accuracy rate decreases as STAI-state score increases both (A) before the mood induction as well as (B) after the mood induction. STAI-state score did not influence RT either (A) before the mood induction or (B) after the mood induction. Each dot represents a single participant.

*9.2.3 Effect of task difficulty on task performance*

As expected based on our previous findings (Exp 1), accuracy rate decreased significantly as a function of task difficulty level (F(2.63, 1503.98) = 1731.32, $p$ < .001), where accuracy rate was significantly different in all task difficulty levels (all $p$s < .001) (Figure 5A). Moreover, RT increased significantly as a function of task difficulty level (F(2.85, 1442.07) = 3913.64, $p$ < .001). RT at each task difficulty level was significantly different from the others (all $p$s < .001; Figure 5B).

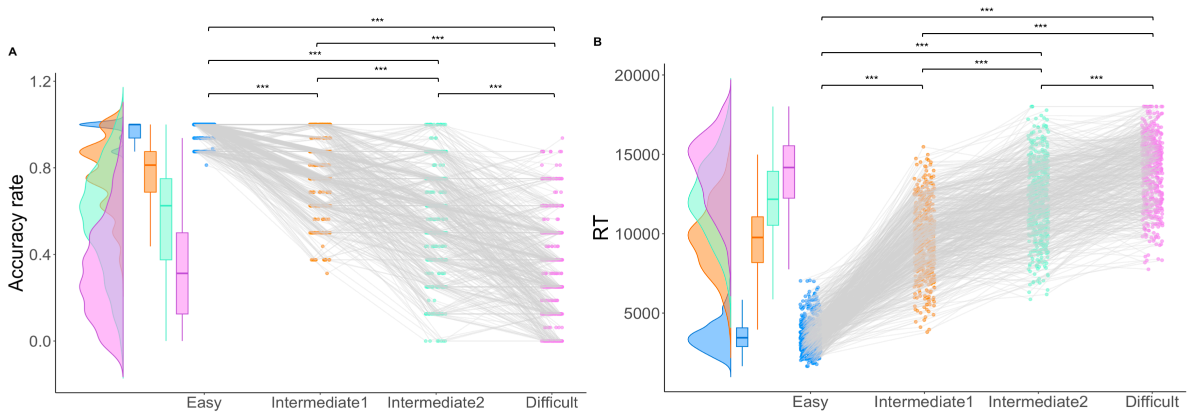


Figure 5. Relationship between difficulty level and effort evaluation. (A) displays linear decrease in accuracy rate with increasing task difficulty. (B) Shows the linear increase in RT as task difficulty increases.

*9.2.4. Effect of task difficulty on flow score and EC scores*

In line with previous findings (Exp 1), our analysis showed that participants rated different subjective flow score based on flow induction task difficulty level ($F(1.77, 660.26) = 52.10$, $p < .001$; Figure 6A). Participants were significantly higher engaged in the two intermediate levels as compared to the easy (both *p*s < .001), and difficult level (both *p*s < .001). There was no significant difference between easy and difficult ($p = .114$). Participants reported significantly higher engagement in the intermediate1 level versus intermediate 2 ($p < .001$).

Moreover, our results showed a significantly different EC score across task difficulty levels ($F(2.57, 956.44) = 14.90$, $p < .001$) (Figure 6B). This linear effect is driven by significantly lower E scores in the easy level compared to the intermediate1 level ($p = .041$), the intermediate2 level ($p = .001$) and the difficult level ($p < .001$), and lower intermediate1 and intermediate2 scores compared to the difficult level (both *p*s < .001) (Figure 6B). No significant effect was found for intermediate1 versus intermediate2 ($p = 1.000$). This again confirms the previous findings that willingness to exert effort in a task decreases as the difficulty increases.

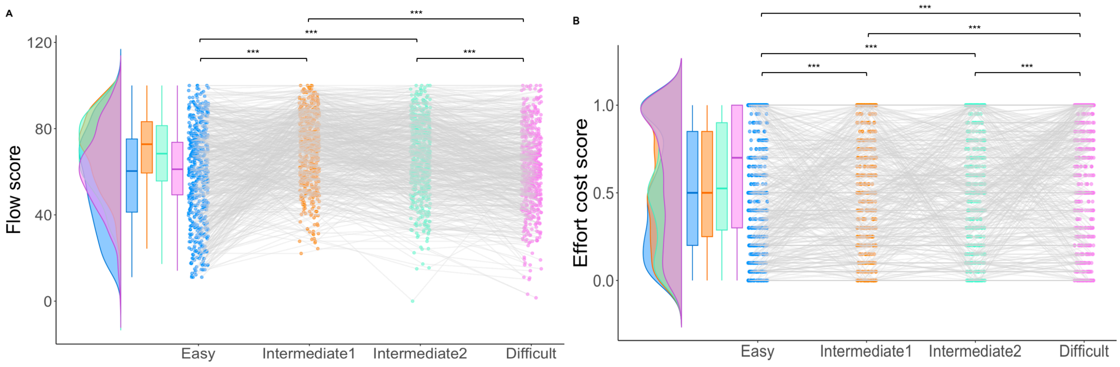


Figure 6. Relationship between difficulty level and effort evaluation.

(A) Visualizing the quadratic effect of higher subjective flow score for easy and difficult tasks.

(B) Effort cost shows a positive increase as the task difficulty increases.

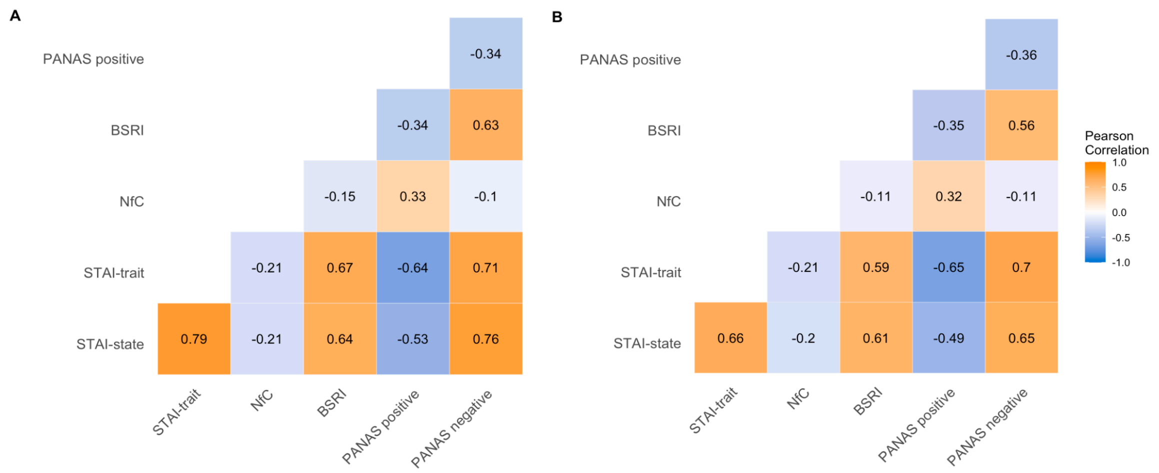


Figure 7. Heatmap showing the correlation coefficients per inventory of A) pre induction score, B) post induction score

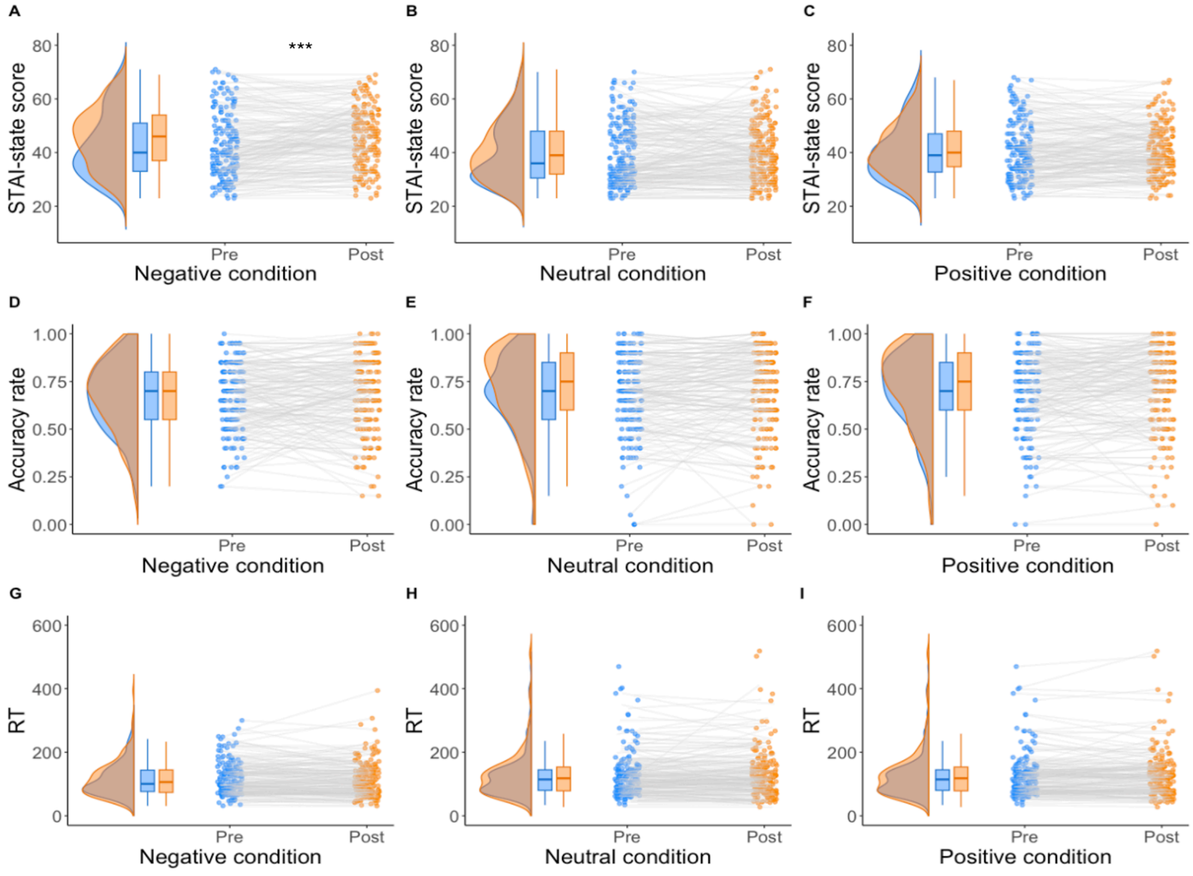


Figure 8. Anxiety and task performance during the SART

(A) STAI-state scores are higher after the mood induction compared to before. There is no significant change in STAI-state score after the mood induction in the neutral (B) and positive condition (C). The accuracy rate does not significantly change after the mood induction in the negative (D), neutral (E), or positive (F) conditions. Similarly, there is no significant difference in RT between before and after the mood induction in the negative (G), neutral (H), or positive (I) conditions. Each dot represents a single participant.

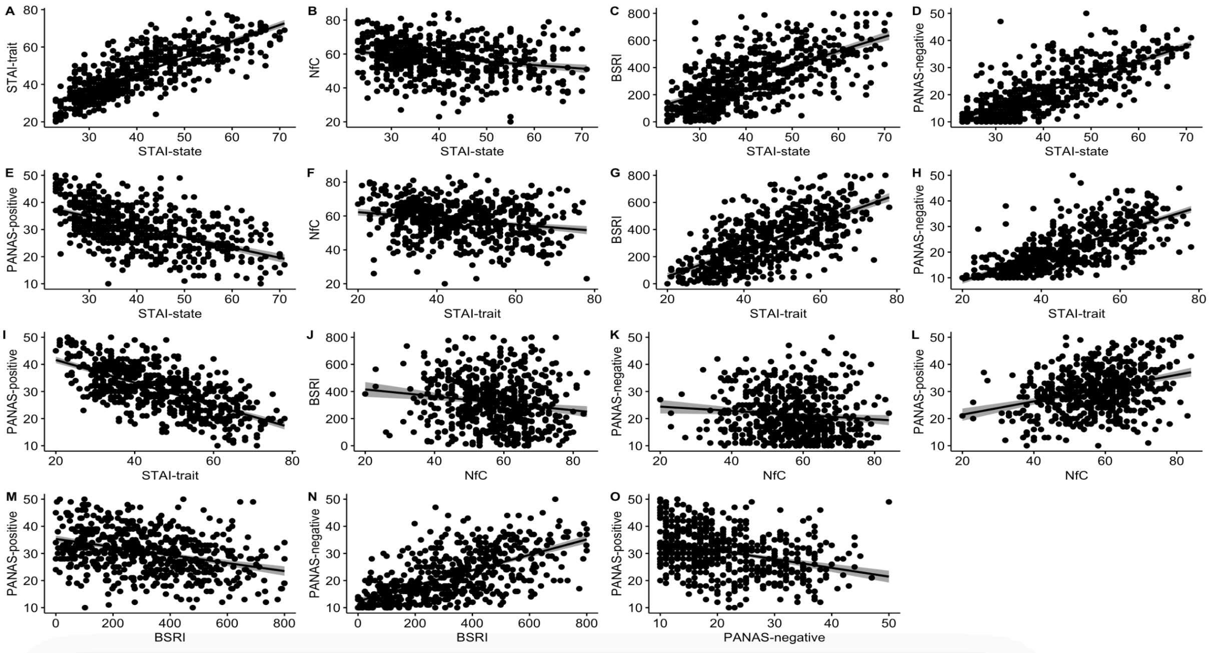


Figure 9. Scatterplots of relationship between the inventories (pre scores). Each dot represents a single participant

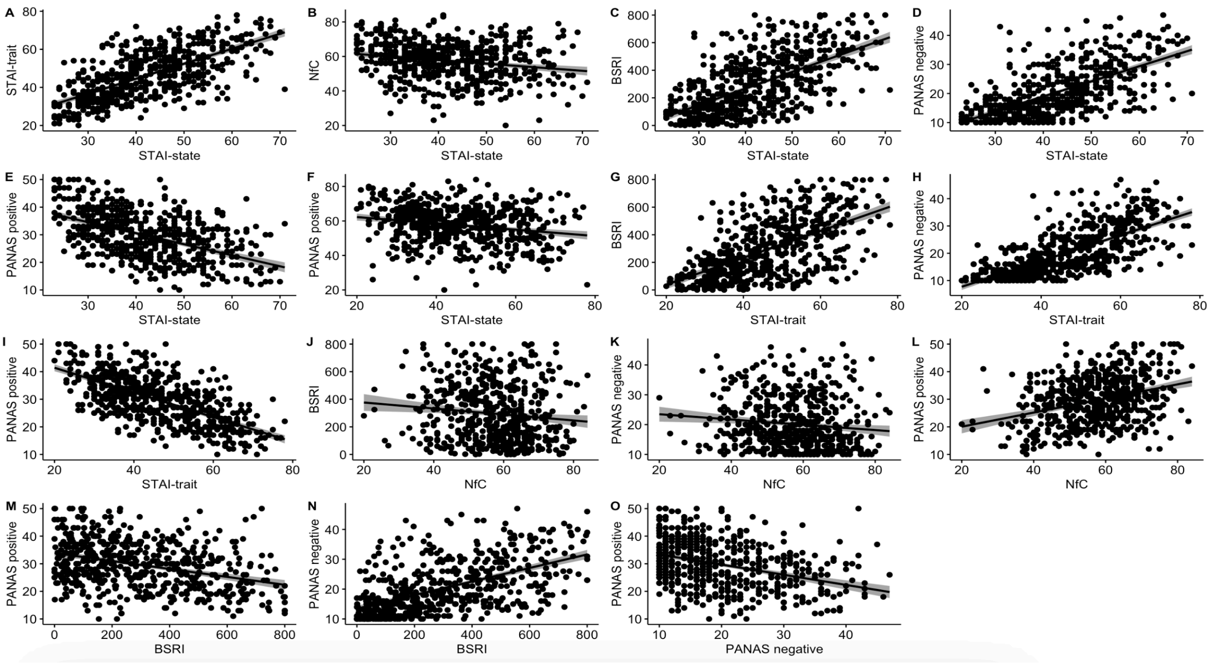


Figure 10. Scatterplots of relationship between the inventories (post scores). Each dot represents a single participant

| Predictors | Estimate | SD | T value | P value | Df |
|---|---|---|---|---|---|
| Difficulty level | -.21 | < .01 | -71.51 | < .001 | 1713 |
| Condition | < .01; - < .01 | .01; .01 | .45; -.45 | .868 | 571 |
| STAI-trait | < .01 | < .01 | .54 | .587 | 571 |
| Difficulty level x condition | < .01; - .01 | < .01; < .01 | .83; -1.58 | .287 | 1713 |
| Difficulty level x STAI-trait | < .01 | < .01 | .75 | .453 | 1713 |
| Condition x STAI-trait | < -.01; < .01 | < .01; < .01 | -.27; .21 | .961 | 571 |
| Difficulty level x condition x STAI-trait | < .01; < -.01 | < .01; < .01 | .20; -.38 | .931 | 571 |
| Covariate years of education | < .01 | < .01 | 1.30 | .193 | 571 |

Table 11. Accuracy rate model output: condition

| Predictors | Estimate | SD | T value | P value | Df |
|---|---|---|---|---|---|
| Difficulty level | .45 | < .01 | 72.20 | < .001 | 1 |
| Condition | < .01; - .01 | .01; .01 | .24; -.96 | .535 | 2 |
| STAI-trait | < -.01 | .01 | -1.75 | .106 | 1 |
| Difficulty level x condition | < - .01; .01 | .01; .01 | .92; .73 | .945 | 2 |
| Difficulty level x STAI-trait | < -.01 | < .01 | -1.04 | .307 | 1 |
| Condition x STAI-trait | < -.01; < .01 | < .01; < .01 | -.56; .64 | .791 | 2 |
| Difficulty level x condition x STAI-trait | < -.01; < .01 | < .01; < .01 | -.13; .56 | .845 | 2 |

Table 12. RT model output: condition

| Predictors | Estimate | SD | T value | P value | Df |
|---|---|---|---|---|---|
| Difficulty level | .54 | .40 | 1.34 | .180 | 571 |
| Condition | -.22; 1.27 | .74;.73 | -.30; 1.74 | .178 | 571 |
| STAI-trait | - .21 | .04 | -4.94 | < .001 | 571 |
| Difficulty level x condition | .09; - .15 | .57; .56 | .15; -.26 | .966 | 571 |
| Difficulty level x STAI-trait | -.07 | .03 | -2.17 | .031 | 571 |
| Condition x STAI-trait | -.04; 0.11 | .06; .06 | -.74; 1.96 | .022 | 571 |
| Difficulty level x STAI-state x STAI-trait | -.04; -.02 | .05; .05 | -.88; -.37 | .44§ | 571 |

Table 13. Flow model output: condition

| Predictors | Estimate | SD | T value | P value | Df |
|---|---|---|---|---|---|
| Difficulty level | .03 | .01 | 6.17 | <.001 | 571 |
| Condition | -.01; -.02 | .02; .02 | -.90; -1.48 | .058 | 571 |
| STAI-trait | < -.01 | < .01 | -1.42 | .155 | 571 |
| Difficulty level x condition | -.10; < -.01 | .01; .01 | -1.63; -.53 | .082 | 571 |
| Difficulty level x STAI-trait | < -.01 | < .01 | -.07 | .95 | 571 |
| Condition x STAI-trait | < .01; < -.01 | < .01; <.01 | 1.67; -.50 | .08 | 571 |
| Difficulty level x STAI-state x STAI-trait | < .01; < -.10 | < .01; <.01 | .21; -.28 | .958 | 571 |
| Covariate age | < .01 | < .01 | .56 | .573 | 571 |

Table 14. Effort cost model output: condition

| Predictors | Estimate | SD | t value | P value | Df |
|---|---|---|---|---|---|
| Involvement: STAI-state x STAI-trait | .01 | .01 | 2.34 | .020 | 572 |
| Involvement: Difficulty level x STAI-state x STAI-trait | < -.01 | <. .01 | -1.04 | .301 | 572 |
| Liking: STAI-state x STAI-trait | .01 | .01 | 1.16 | .246 | 571 |
| Liking: Difficulty level x STAI-state x STAI-trait | < .01 | < .01 | -.85 | .841 | 571 |
| Ability: STAI-state x STAI-trait | 01 | < .01 | 2.13 | .034 | 565 |
| Ability: Difficulty level x STAI-state x STAI-trait | < .01 | < .01 | .09 | .927 | 572 |
| Time: STAI-state x STAI-trait | .01 | .01 | 1.98 | .048 | 572 |
| Time: Difficulty level x STAI-state x STAI-trait | < .01 | < .01 | 0.28 | .780 | 572 |

Table 15. Flow subcomponent model output: relevant interaction effects

| Predictors | Estimate | SD | t value | P value | Df |
|---|---|---|---|---|---|
| Involvement: STAI-state for low STAI-trait | -.54 | .10 | -5.48 | < .001 | 299 |
| Involvement: STAI-state for high STAI-trait | -.27 | .11 | -2.52 | .005 | 273 |
| Ability: STAI-state for low STAI-trait | -.34 | .11 | -3.21 | .001 | 299 |
| Ability: STAI-state for high STAI-trait | .02 | .11 | .19 | .626 | 273 |

Table 16. Flow subcomponent model output: significant interaction effects

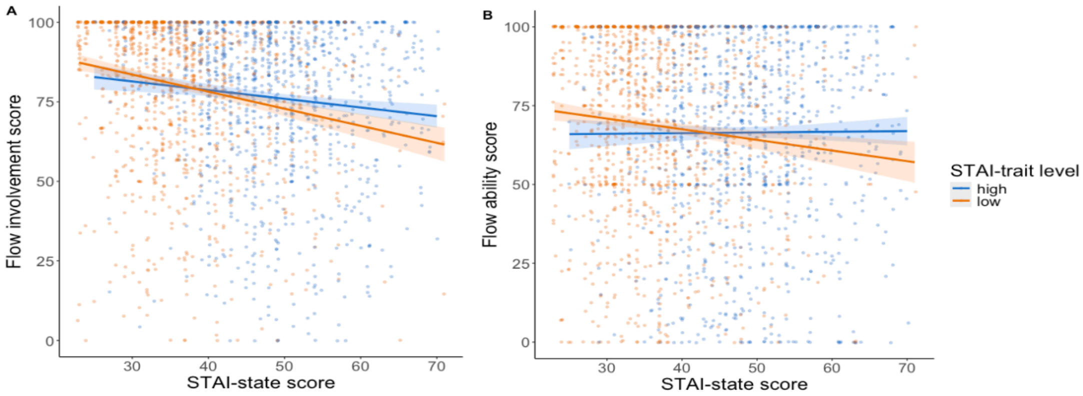


Figure 11. The effect of anxiety of A) flow subcomponent involvement, B) flow subcomponent ability.
